# Doc2b Ca^2+^-binding site mutants act as a gain of function at rest and loss of function during neuronal activity

**DOI:** 10.1101/536581

**Authors:** Quentin Bourgeois-Jaarsma, Matthijs Verhage, Alexander J. Groffen

## Abstract

Communication between neurons involves presynaptic neurotransmitter release which can be evoked by action potentials or occur spontaneously as a result of stochastic vesicle fusion. The Ca^2+^-binding double C_2_ proteins Doc2a and –b regulate both spontaneous and asynchronous evoked release, but the mechanism remains unclear. Here, we compared wildtype Doc2b with two Ca^2+^ binding site mutants named DN and 6A, respectively considered gain-and loss-of function mutants and carrying the substitutions D218,220N or D163,218,220,303,357,359A. We found that both mutants bound phospholipids at low free Ca^2+^ concentrations and were membrane-associated in neurons at rest, mimicking a Ca^2+^ activated state. Their overexpression in hippocampal primary neurons culture had similar effects on spontaneous and evoked release, inducing higher mEPSC frequencies and increased short-term depression. Together, these data suggest that the DN and 6A mutants both act as gain-of-function mutants at resting conditions but as loss-of-function during neuronal activity.

## Introduction

Regulated exocytosis is strictly dependent on *Soluble N-ethylmaleimide-sensitive-factor Attachment protein REceptor* (SNARE) proteins, Ca^2+^-sensors and a number of accessory proteins (Südhof, 2013). Neurotransmitter release is either triggered by action potentials (APs) (Südhof, 2004, 2013; Chapman, 2008; Kaeser and Regehr, 2014; Meriney et al., 2014) or occurs spontaneously at resting membrane potential (Fatt and Katz, 1950; Kaeser and Regehr, 2014).

Evoked release consists of synchronous and asynchronous release components (Otsu et al., 2004; Chapman, 2008; Bacaj et al., 2013). Fast, synchronous release triggered by local Ca^2+^influx (nano & micro-domain) occurs in less than a millisecond (Kaeser and Regehr, 2014; Neher, 2015) and is governed by the fast Ca^2+^ sensors Syt-1, 2 or 9 (Xu et al., 2007). Another class of high affinity Ca^2+^ sensors with slow kinetics such as Syt-7 mediates asynchronous release (Saraswati et al., 2007; Xue et al., 2010; Bacaj et al., 2013; Luo and Südhof, 2017). In synapses lacking the fast sensor, diminished synchronous release is accompanied by increased asynchronous release as shown for Syt-1 (Geppert et al., 1994; Maximov and Südhof, 2005) and Syt-2 (Nagy et al., 2006; Pang et al., 2006; Sun et al., 2007).

Unlike evoked release, spontaneous release is AP-independent and occurs as a stochastic process with a probability that appears to be regulated by intracellular Ca^2+^ (Xu et al., 2009; Groffen et al., 2010; Ermolyuk et al., 2013). Spontaneous release is important for nervous system functioning as it is involved in synapse maturation, maintenance and synaptic plasticity (McKinney et al., 1999; Sutton et al., 2004; Ehlers et al., 2007; Lee et al., 2010). Like asynchronous release, its frequency is suppressed by Syt-1 and Syt2 (Xu et al., 2009; Kochubey and Schneggenburger, 2011; Courtney et al., 2018) and stimulated by double C_2_(Doc2) proteins (Groffen et al., 2010; Pang et al., 2011; Williams and Smith, 2018).

Doc2a, -b and –c isoforms together constitute the Doc2 protein family. Unlike Doc2a and –b, the third isoform does not have functionally conserved C_2_ domains. Doc2a is mainly expressed in the adult brain while Doc2b is more widely expressed in the nervous system and various neuroendocrine tissues (Verhage et al., 1997; Korteweg et al., 2000). Both Doc2a and –b contribute to spontaneous release as shown in knockout and knock-down models (Groffen et al., 2010; Pang et al., 2011). A recent study revealed that they regulate spontaneous excitatory and inhibitory activity, respectively (Courtney et al., 2018). In cell-free assays, Doc2b interacts with the SNARE complex to promote fusion of SNARE-liposomes (Groffen et al., 2010; Yao et al., 2011). Doc2b C_2_ domains bind to phosphatidylserine-containing membranes in a Ca^2+^-dependent manner (Groffen et al., 2010) but also to PI(4,5)P_2_, a phospholipid enriched on the cytoplasmic leaflet of the plasma membrane (Michaeli et al., 2017).

The N-terminal domain of Doc2a/b interacts with Munc13 via a Munc13 interacting domain (MID; Figure 1A, B) in HEK293 cells (Duncan et al., 1999), PC12 cells (Orita et al., 1997) chromaffin cells (Friedrich et al., 2010) and neurons (Hori et al., 1999; Xue et al., 2018). This interaction is sufficient for co-translocation of Munc13 together with Doc2 upon phorbol ester (a DAG homologue) stimulation (Duncan et al., 1999; Hori et al., 1999; Groffen et al., 2004). Phorbol esters potentiate exocytosis in a Ca^2+^-independent way relying on Doc2/Munc-13 interaction (Hori et al., 1999; Duncan et al., 2000; Groffen et al., 2004). Consistently, Doc2 overexpression causes a Ca^2+^-independent, Munc-13 dependent release increase upon phorbol ester stimulation (Friedrich et al., 2013) and conversely, blockade of the Doc2-Munc13 interaction by synthetic peptides abolishes phorbol ester potentiation (Hori et al., 1999). Munc13-1 is necessary for the Doc2b-induced priming of secretory granules in chromaffin cells (Houy et al., 2017). However, alteration of this interaction by mutations in the MID domain have no effect on Ca^2+^-induced Doc2b migration to the membrane (Groffen et al., 2004; Gaffaney et al., 2014). The presence of PI(4,5)P_2_ targets Doc2b to the plasma membrane upon [Ca^2+^]_i_elevation (Michaeli et al., 2017). Hence, Doc2b could support exocytosis by both of several mechanisms: i) together with Munc-13 for vesicle priming or superpriming; ii) Ca^2+^-dependently by enhancing membrane fusion.

**Figure 1.**
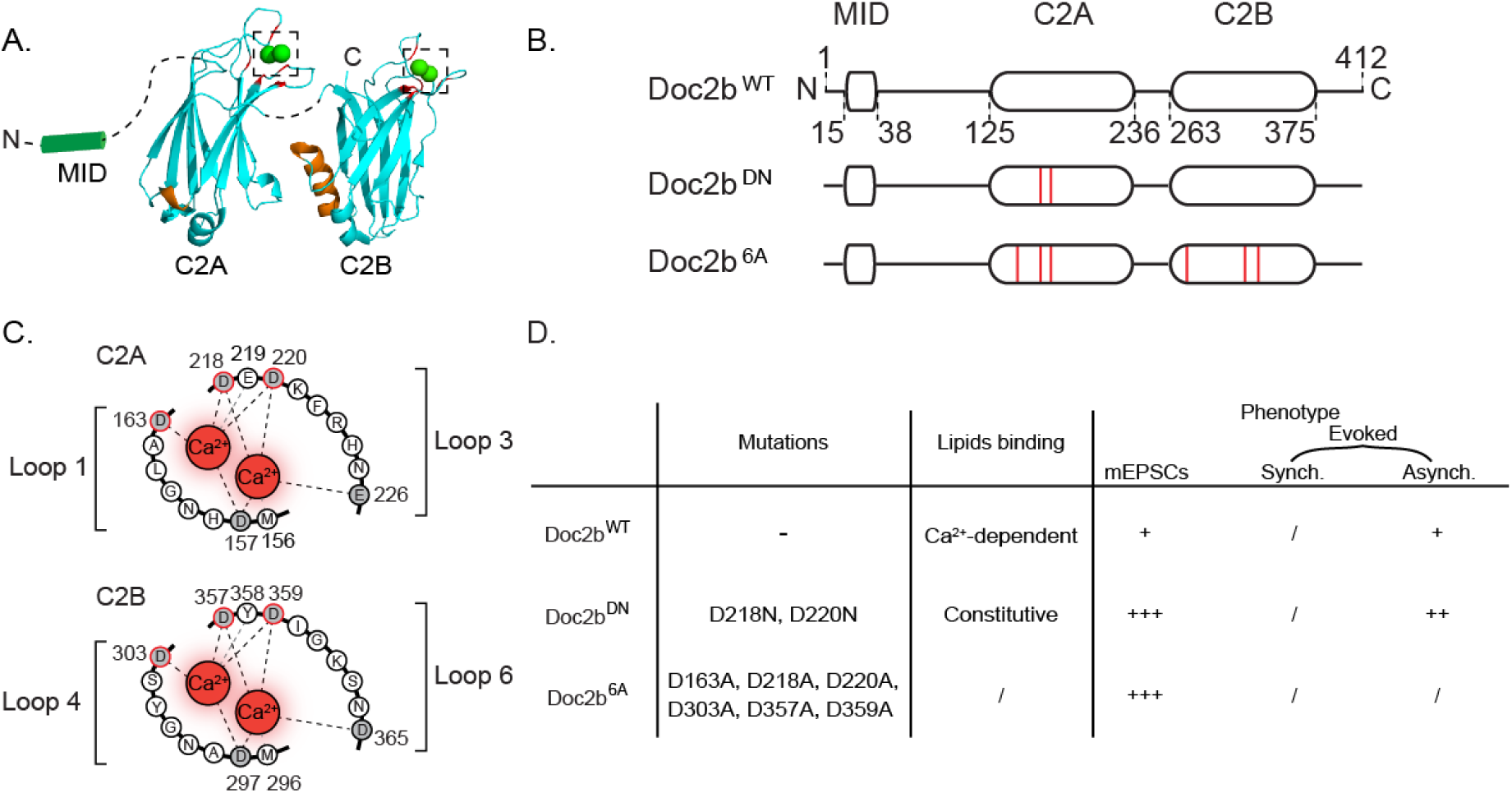
Molecular and phenotypic properties of Doc2b and its Ca^2+^-binding site mutants. (**A**) Cartoon showing C_2_ domain structures of Doc2b based on crystallography (Giladi et al., 2013). Aspartates involved in Ca^2+^binding are marked in red; poly-lysine sequences for SNARE complex interaction are marked in orange (Rigsby and Parker, 2016). Dotted lines represent linker sequences between domains. Dotted squares highlight Ca^2+^-binding pockets enlarged in C. (**B**) Linear representation of Doc2b ^WT^ and two previously investigated mutants both mutants Doc2b^DN^ and Doc2b^6A^(red lines indicate amino acid substitutions). (**C**) Aspartates substituted in Doc2b^DN^ (D218, 220N) or Doc2b^6A^(D163, 218, 220, 303, 357, 359A in). (**D**). Summary of functional effects of Doc2b^DN^and Doc2b^6A^mutations as measured by phospholipid binding in cell-free systems or by electrophysiology in cultured neurons (Groffen et al., 2010; Pang et al., 2011).

Doc2a/b have a high Ca^2+^ affinity with half-maximal membrane binding at 450 nM and 175 nM respectively in chromaffin cells (Groffen et al., 2006). Ca^2+^ binding onto their C_2_ domains requires five acidic aspartate amino acids (Figure 1B) in close proximity to hydrophobic loops which interact with the membrane after Ca^2+^activation, resulting in reversible translocation (Groffen et al., 2004, 2010). A similar mechanism occurs in Syt-1 (Frazier et al., 2003), except that Syt-1 is anchored to synaptic vesicles by a N-terminal transmembrane domain and has a lower apparent Ca^2+^ affinity. Doc2b and Syt-1 compete for SNARE protein binding (Groffen et al., 2010; Yao et al., 2011), which suggest a partially shared mechanism in Ca^2+^-secretion coupling as recently mentioned for spontaneous release (Courtney et al., 2018).

It is still debated whether Doc2b acts as a direct Ca^2+^-sensor or as a structural element supporting Ca^2+^-dependent secretion by another process. Neutralization of two critical aspartates D218 and D220 in the C_2_A domain of Doc2b (Figure 1B-D) induce Ca^2+^-independent membrane-binding of the domain (Friedrich et al., 2008; Xue et al., 2015). This mutant, designated Doc2b^DN^, was therefore considered a gain-of-function mutant. When Doc2b^DN^ expression caused a rise of the spontaneous release rate (mEPSCs), this was taken to support a role as Ca^2+^ sensor (Groffen et al., 2006, 2010). Another Ca^2+^-ligand mutant designated Doc2b^6A^, in which six aspartates were substituted by alanines (Figure 1B-D), is unable to bind Ca^2+^and therefore considered a loss-of-function mutant (Pang et al., 2011). This mutant still rescues spontaneous release in Doc2 double knock-down neurons, suggesting a Ca^2+^-independent mechanism. Nevertheless, a recent study suggested that this mutant may enhance spontaneous release by mimicking a Ca^2+^-bound state (Courtney et al., 2018).

As another point of debate, several studies reported reduced asynchronous release in Doc2a knock-out neurons which is rescued by Doc2b expression (Gaffaney et al., 2014; Xue, Gaffaney, & Chapman, 2015; Yao et al., 2011; Figure 1D) and further enhanced by expression of Doc2b mutants, leaving room for debate whether Doc2b acts selectively on spontaneous release, selectively on asynchronous release, or contributes to both processes.

Here we directly compared both mutants and wildtype (WT) Doc2b for their Ca^2+^-dependent membrane-binding activity, their subcellular localization and their effects on spontaneous and evoked neurotransmission. Under resting conditions both mutants showed a gain-of-function behavior. At high Ca^2+^ however, both mutants caused significantly higher short-term depression, suggesting a loss of function. These results reconcile previous findings and show that Doc2b can affect both spontaneous release and short-term plasticity.

## Results

### Aspartate substitutions in Doc2b^DN^and Doc2b^6A^cause constitutive membrane association in resting neurons

Activity-dependent plasma membrane binding is an established feature of Doc2b (Groffen et al., 2006). We first tested the Ca^2+^-dependent membrane binding of the Doc2b^DN^and Doc2b^6A^mutants, previously described to be gain-and loss-of function variants respectively (Groffen et al., 2010; Pang et al., 2011) under resting conditions (Figure 2A) and during stimulation (Figure 2B-C). In samples fixed at resting conditions, eGFP-tagged Doc2b^WT^was localized homogeneously throughout the cytosol similar to soluble eGFP, as expected for a cytosolic protein (Figure 2A) and in line with previous findings (Groffen et al., 2006). In contrast, eGFP-tagged Doc2b^DN^and Doc2b^6A^both showed plasma membrane enrichment (Figure 2A), again consistent with previous studies (Friedrich et al., 2008; Houy et al., 2017).

**Figure 2.**
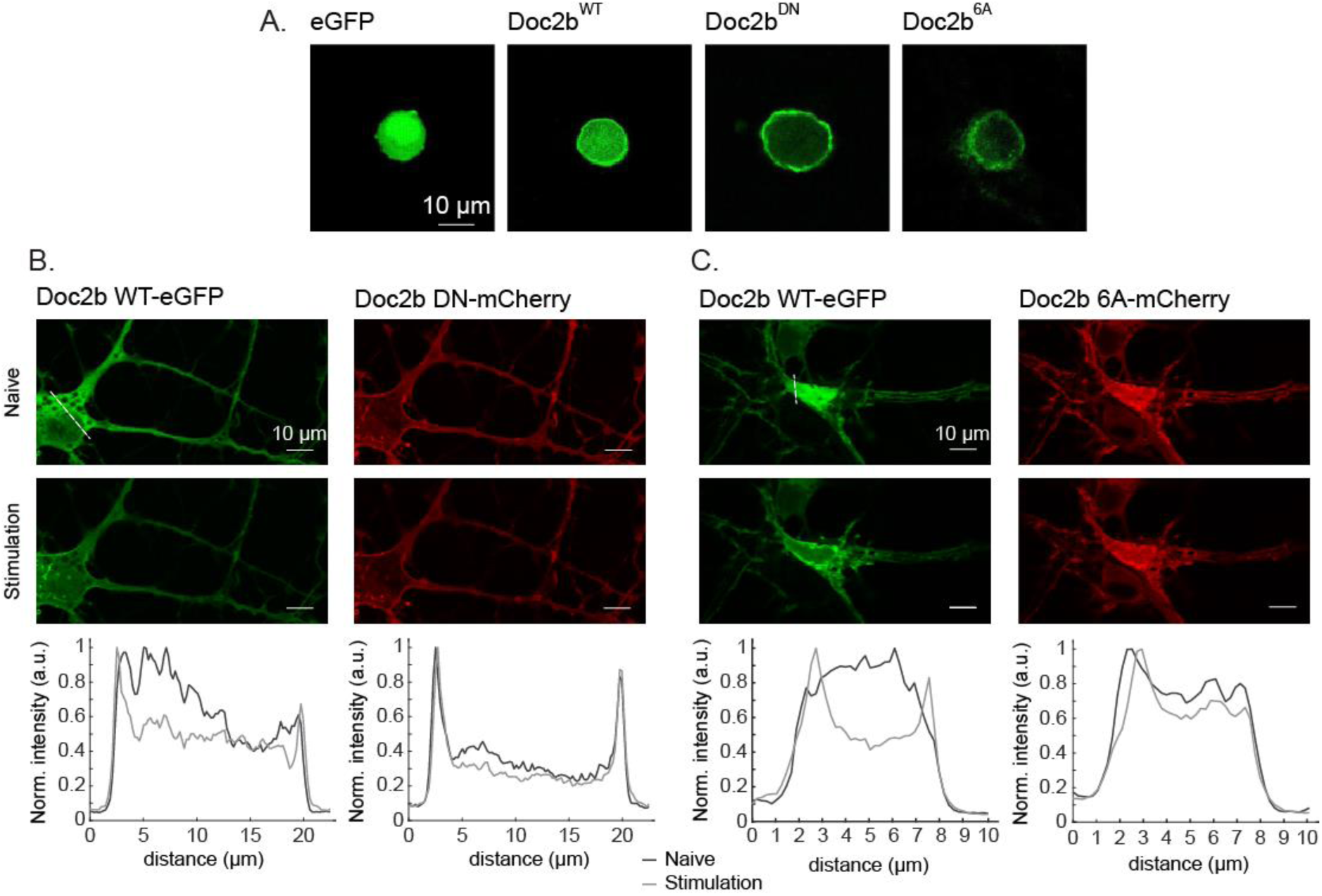
Ca^2+^binding site mutants Doc2b^DN^and Doc2b^6A^show increased plasma membrane binding under resting conditions. (**A**) Confocal microscopy on fixed cells and (**B-C**) live epifluorescence microscopy from autaptic neurons overexpressing Doc2B wildtype and mutants before and during 40Hz stimulation for 15 seconds. Images represent an average from 5 frames for each condition. Signal intensity comparisons (lower panels in B-C) correspond to line scan profiles reported by dotted white lines (upper panels in B-C).

To assess the activity dependence of membrane association, we performed confocal live imaging upon field stimulation at 40Hz for 15 s (Figure 2B). To control for cell-to-cell differences, mCherry-tagged mutants were co-expressed with eGFP-tagged Doc2b^WT^in the same neurons. Again, both Doc2b^DN^and Doc2b^6A^were membrane-bound while Doc2b^WT^showed a homogeneous cytoplasmic localization at rest. During repetitive stimulation, Doc2b^WT^showed a clear plasma membrane association, whereas the distribution of both mutants showed only a small change (Figure 2B, C). Doc2b^DN^slightly enhanced its membrane localization during repetitive stimulation, while Doc2b^6A^revealed a slight decrease in plasma membrane binding (Figure 2B, C). Thus, in living neurons, both mutants exhibit increased plasma membrane binding at rest while activity-induced plasma membrane binding is impaired.

### Phospholipid-binding properties of Doc2b mutants

The altered subcellular localization of Doc2b^DN^and Doc2b^6A^could result from altered C_2_-phospholipid interactions. To investigate this, wildtype and mutant C_2_A and C_2_AB fragments were expressed as recombinant proteins in bacteria. The mutants were expressed to the expected molecular mass as verified by SDS-PAGE (Figure 3A, I). C_2_-phospholipid binding was measured in a liposome aggregation assay (Connell, Scott, & Davletov, 2008), using calibrated EGTA-buffered Ca^2+^solutions and liposomes composed of 25% DOPS and 75% DOPC (Friedrich et al., 2008). To test phospholipid binding by the C_2_A domain, a glutathione-S-transferase tag was used to induce self-dimerization, so that phospholipid binding causes liposome clustering which can be measured as an absorbance increase at 350 nm. Addition of a GST-C_2_A^WT^protein fragment to a Ca^2+^-containing solution caused rapid liposome clustering (Figure 3B, E). This activity was strictly Ca^2+^-and protein-dependent and followed a sigmoid dose dependence giving in EC_50_ of 435 ± 31 nM in line with previous reports (Friedrich et al., 2008). GST-C_2_A^WT^remained membrane-bound at high free Ca^2+^concentrations in the 1-10 µM range. In contrast, both GST-C_2_A^DN^and GST-C_2_A^3A^(here called 3A because only 3 of the total 6 aspartates reside in the C2A domain) showed strong phospholipid binding at low free Ca^2+^concentrations in the range 0 - 500 nM. At higher [Ca^2+^]_free_, both mutants displayed a loss of phospholipid association (Figure3 C-D).

**Figure 3.**
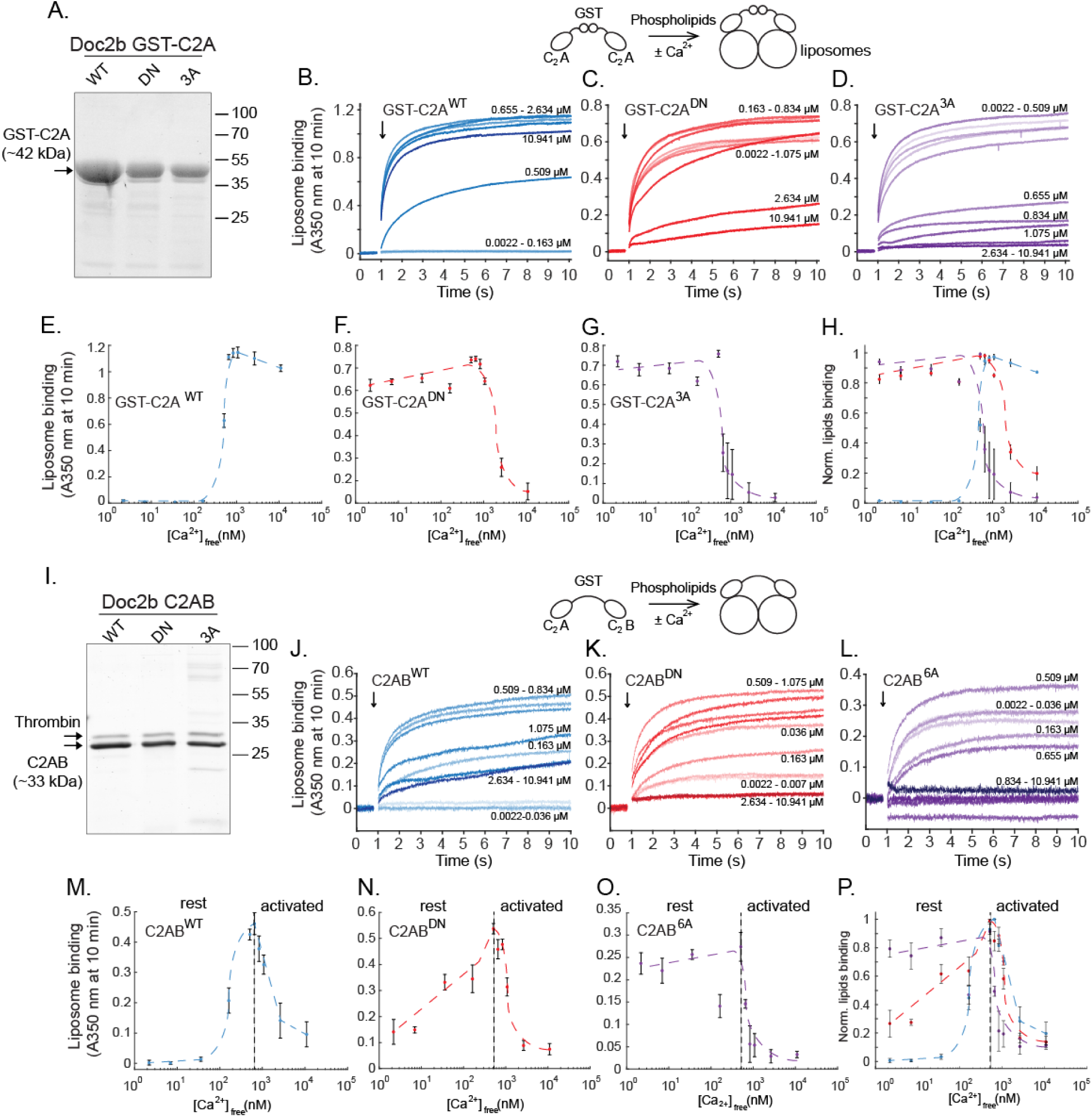
Ca^2+^binding site mutations cause constitutive phospholipid binding of C_2_A (A-H) and C_2_AB fragments (I-P) of Doc2b. (**A**) Recombinant GST-C_2_A fusion protein of wildtype and mutants Doc2b visualized by SDS-PAGE and Sypro Ruby staining. (**B-D**). The addition of GST-C_2_A fragments (arrows) to 75 % DOPC 25 % DOPS-containing liposomes caused liposome aggregation. Kinetic measurements of absorbance at λ=350 nm were performed in various free Ca^2+^concentrations in calibrated Ca^2+^/EGTA solutions. (**E-H**) Ca^2+^-dependence of phospholipid binding by GST-C_2_A^WT^, GST-C_2_A^DN^and GST-C_2_A^3A^. (**I**) Recombinant C_2_AB fragments of Doc2b^WT^, Doc2b^DN^, and Doc2b^6A^. (**J-L**) Kinetic absorbance measurements of 75 % DOPC/25 % DOPS liposomes. (M-P) Ca^2+^-dependence of liposome aggregation activity for each C_2_AB construct. At low (<50 nM) [Ca^2+^]_free_, both mutants showed increased phospholipid binding activity while the wildtype constructs did not function. However, in high free Ca^2+^concentrations condition (>1 µM), a strong reduction in activity was observed for all constructs except C_2_A^WT^. Data are represented as mean ±SEM from N=5 and N=4 independent measurements for GST-C_2_A and C_2_AB recombinant fragments respectively. Dashed lines indicate manually drawn trendlines used as visual aids.

To test phospholipid aggregation by C_2_AB fragments, the GST tag was removed using thrombin cleavage. Liposome clustering by C_2_AB^WT^increased Ca^2+^-dependently to reach a half-maximum at 176 ± 38 nM (Figure 3M) consistent with reported data (Groffen et al., 2006; Giladi et al., 2013; Gaffaney et al., 2014). A maximum occurred at approximately 700 nM,followed at higher [Ca^2+^]_free_ by a strong decrease in the absorbance signal. At the lowest tested [Ca^2+^]_free_of 2.2 nM, the C_2_AB^DN^fragment already showed partial activity (Figure 3N). This activity increased with higher [Ca^2+^]_free_This increase presumably reflects Ca^2+^-dependent activity from the intact C_2_B domain. At high Ca^2+^concentrations above 500 nM, a decrease was observed reminiscent of the data obtained with C_2_AB^WT^. The C_2_AB^6A^fragment showed near-complete liposome binding at the lowest [Ca^2+^]_free_and no prominent Ca^2+^dependency in the 0 – 500 nM range of [Ca^2+^]_free_(‘rest’, Figure 3O), followed again by a signal decrease above 500 nM. Taken together, in this cell-free liposome clustering assay, both Doc2b^DN^and Doc2b^6A^behave as a gain-of-function mutants at low [Ca^2+^], but as loss-of-function mutants at high [Ca^2^].

### Doc2b^DN^and ^6A^overexpression similarly enhances spontaneous release

To measure the effect of Doc2b on synaptic activity, wildtype hippocampal neurons were cultured either in networks (Figure 4A-C) or on glial micro-islands to promote self-innervation (autaptic neurons; Figure 4D-F). In networks, spontaneous release was measured in presence of 1 µM tetrodotoxin (TTX) to block voltage-gated sodium channels. Doc2b was expressed by lentiviral vectors in wildtype neurons. Western-blot confirmed low expression levels of endogenous Doc2b (Korteweg et al., 2000) which were detectable in lysate from brain and cultured cortical neurons (Figure S1C). Viral infection induced Doc2b protein levels much higher than endogenous levels (Figure S1A).

**Figure 4.**
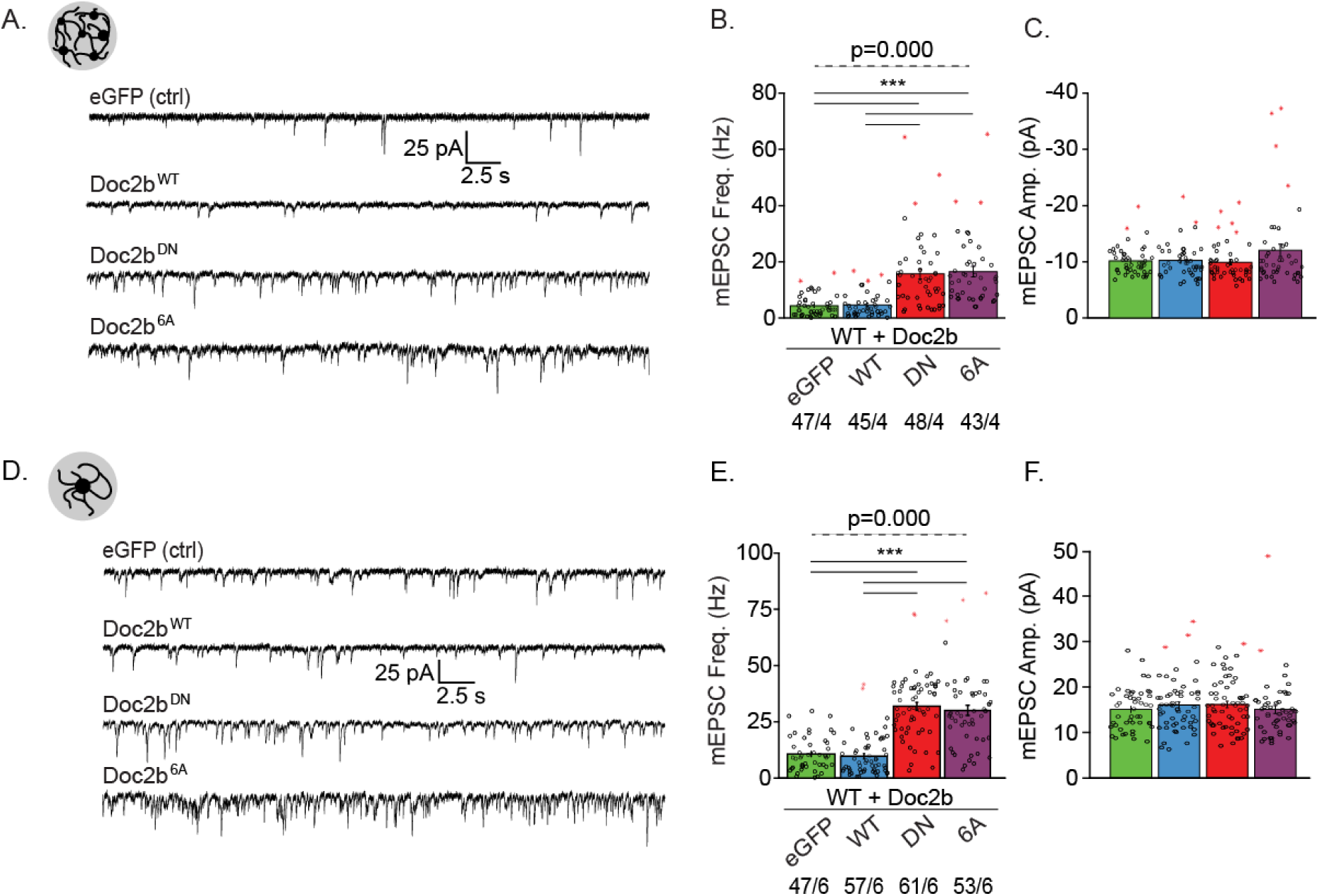
Doc2b^DN^and Doc2b^6A^mutations enhance the frequency of spontaneous release in wildtype hippocampal neurons. (**A**) Representative mEPSC recordings in wildtype neurons overexpressing eGFP (ctrl), Doc2b wildtype (Doc2b^WT^) and mutants (Doc2b^DN^, Doc2b^6A^), cultured in networks in presence of 1 µM TTX to block action potential and 20 µM gabazine. (**B**) Quantification of spontaneous neurotransmitter release frequency and (**C**) amplitude. (**D**) Representative recordings in isolated neurons cultured on microglial islands (autapses). (**E-F**) Quantification. Data are represented as mean ±SEM; n, number of measurements, N, number of independent experiments. Kruskall Wallis ANOVA, Pairwise Post-hoc tests (***, p<0.005).). Red stars signs indicate outliers.

In both networks and autapses, overexpression of wildtype Doc2b did not change the frequency of miniature excitatory postsynaptic currents (mEPSCs), consistent with previous observations (Groffen et al., 2010). In contrast, overexpression of Doc2b^DN^or Doc2b^6A^induced an approximately 3-fold increase of the mEPSC frequency (Figure 4B,E; see Supplemental Table 1 for statistical tests). In all cases, the mEPSC amplitude, charge and rise time were not changed by Doc2b overexpression.

In autapses, a small but significant effect on mEPSC decay time was observed in both mutants (Figure S2B, effect size (r): 1.260, Supplemental Table 1). This effect on mEPSC decay was not replicated in continental networks and may be attributable to less accurate fitting of exponential decay curves in groups with extremely high mEPSC frequencies. Taken together, in a direct comparison, the aspartate substitutions in Doc2b^DN^and Doc2b^6A^have similar gain of function effects on the spontaneous release frequency in hippocampal neurons, which confirms previous observations (Groffen et al., 2010; Pang et al., 2011; Courtney et al., 2018). In conclusion, the effect of mutant Doc2b on spontaneous mEPSC frequency parallel the changes in *in vitro* plasma membrane binding and phospholipid clustering (i.e. a gain of function in resting conditions).

### Doc2b^DN^or Doc2b^6A^overexpression alter evoked release

The above effects on spontaneous release prompt the question whether the Doc2b mutants also affect synaptic release and plasticity during neuronal activity. We next recorded EPSCs induced by a single action potential (AP) or paired APs at varying intervals in wildtype autaptic neurons. Overexpression of Doc2b^WT^did not significantly affect the first evoked amplitude and charge but neurons exhibited a slight reduction compared to control cells expressing eGFP alone. In a recent study (Toft-Bertelsen et al., 2016), Doc2b^WT^was shown to disperse syntaxin-1 from plasma membrane clusters, thereby inhibiting Ca^2+^currents through voltage gated Ca^2+^channels (VGCCs) in chromaffin cells. To test if a similar mechanism occurs in neurons, we compared the single EPSC amplitude and subsequent calcimycin-evoked charge transfer in wildtype autaptic neurons expressing either eGFP control or overexpressing Doc2b^WT^. The Ca^2+^ionophore bypasses VGGCs and triggers exocytosis by an artificial Ca^2+^influx. No significant change was observed in the EPSC amplitude, nor in the charge transfer induced by calcimycin (Figure S3A-D), indicating that Doc2b does not inhibit synaptic strength by modulating Ca^2+^influx.

When expressing mutant Doc2b^DN^or Doc2b^6A^however, the EPSC charge was larger than in cells expressing Doc2b^WT^(Figure 5A-B). This larger EPSC amplitude was associated with a significantly stronger paired pulse depression at intervals from 20 to 1000 ms, with the strongest difference in the 20 – 200 ms range (Figure 5C). Again, the Doc2b^DN^and Doc2b^6A^mutants caused similar changes.

**Figure 5.**
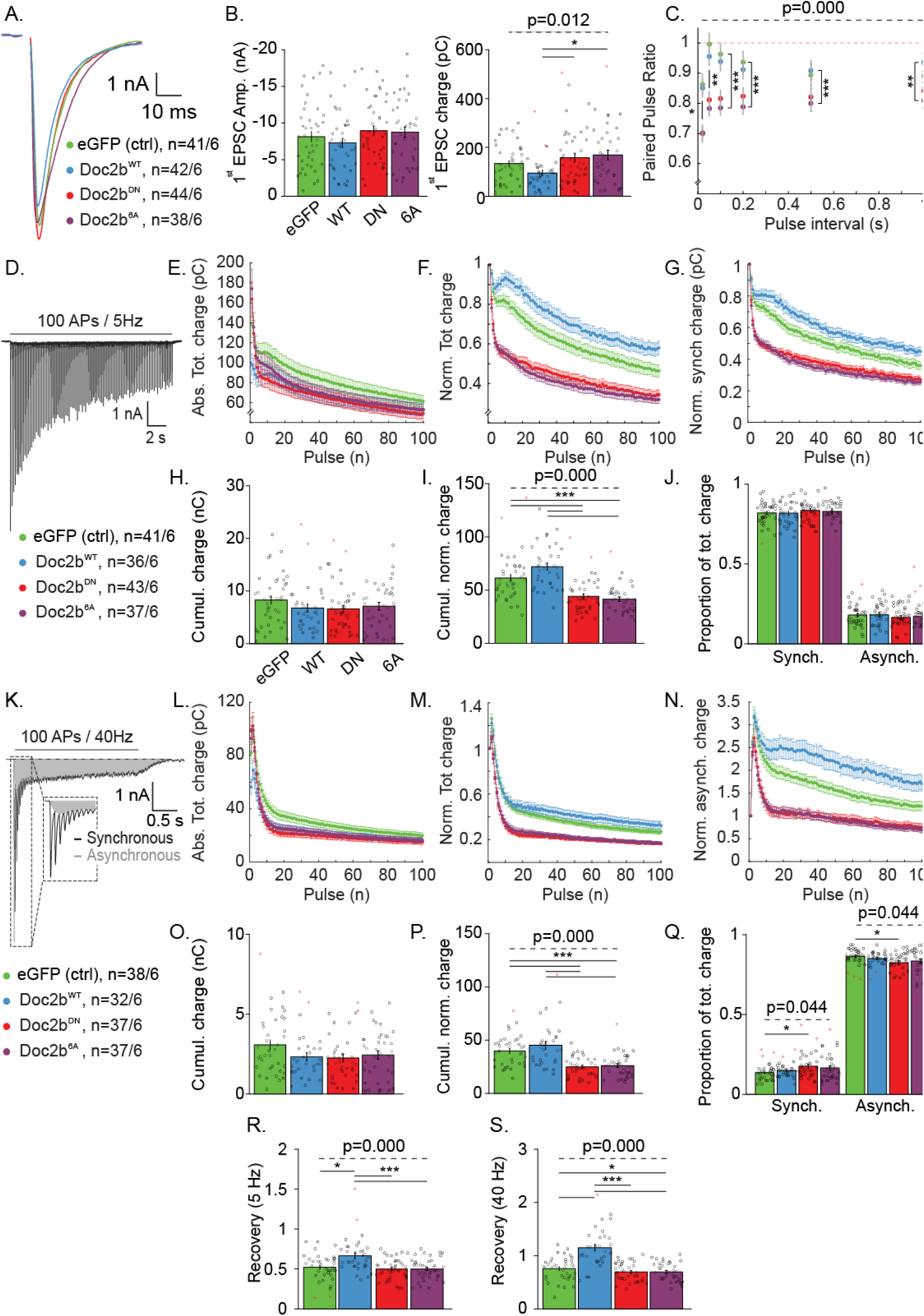
Doc2b mutants affect short term plasticity in autaptic neurons. (**A**) Averaged trace of a single EPSC from wildtype neurons overexpressing Doc2b^WT^and mutants. (**B**) Quantification of 1^st^EPSC charge and (**C**) paired pulse ratio. Doc2b^DN^and Doc2b^6A^affect initial EPSC and short term plasticity. (**D-J**) Rundown during repetitive stimulation at 5 Hz. (**D**) Representative example, (**E**) Absolute and (**F**). Normalized total charge, (**G**) Normalized synchronous release (which is the predominant component of charge transfer at this low stimulation frequency). (**H**) Cumulative total charge and (**I**) Cumulative normalized total charge during the entire train, (**J**) Proportion of the synchronous and asynchronous component. (**K-Q**) Rundown during repetitive stimulation at 40 Hz. (**K**) Representative current in control neurons. (**L**) Absolute and (**M**) Normalized total charge. (**N**) Normalized asynchronous release (which is the predominant component of charge transfer at this high frequency). (**O**) Cumulative absolute and (**P**) cumulative normalized charge during the entire train. (**Q**) Proportion of the synchronous and asynchronous component. (**R**) Recovery, measured as the normalized response charge after 100 stimulations at 5 Hz followed by 2s rest. (**S**) Recovery, measured similarly after a 40 Hz train. Data are represented as mean ±SEM; Kruskall Wallis ANOVA and one way repeated ANOVA, Pairwise Post-hoc tests (*, p<0.05, **, p<0.01, ***, p<0.005). Red stars signs indicate outliers.

To investigate further how sustained evoked release was affected by Doc2b^DN^and Doc2b^6A^, we performed repetitive stimulation with 100 APs at low (5 Hz) and high (40 Hz) frequency (Figure 5D-S). Overexpression of Doc2b^DN^and Doc2b^6A^caused a similar fast depression of the EPSC charge, both at 5 Hz (Figure 5F) and 40 Hz (Figure 5M) yet without altering the total charge transfer (Figure 5H, O). This phenotype was more evident for normalized than for absolute EPSC charges (compare Figure 5F with E and Figure 5M with L respectively), suggesting that the larger initial EPSC charge in mutant expressing neurons contributes to the phenotype. In contrast, overexpression of wildtype Doc2b did not cause any changes in depression compared to GFP-expressing control neurons.

During repetitive stimulation the reduced synchronous release component is accompanied by an asynchronous release component. In view of previous observations implicating Doc2 proteins in asynchronous release (Yao et al., 2011), we tested if Doc2b overexpression affects the proportion of synchronous and asynchronous components to the total EPSC charge (Figure 5J, Q). During repetitive stimulation at 5 Hz, the expression of wildtype or mutant Doc2b did not affect the balance between synchronous and asynchronous release (Figure 5J). Note that the asynchronous component is generally small for low frequency stimulation. Also during 40 Hz stimulation, the balance between synchronous and asynchronous was unaffected by Doc2b^WT^overexpression. However, a small change in favor of synchronous release was seen in Doc2b^DN^expressing neurons (Figure 5Q), but not in Doc2b^6A^-expressing neurons, coherent with the higher EPSC charge (Figure 5C).

To assess the rate of synaptic recovery (Figure 5R and S) we measured the EPSC from a single AP, triggered 2 seconds after the end of a 5 or 40 Hz train. Interestingly, cells expressing wildtype Doc2b showed a higher recovery rate than GFP control cells. This activity was impaired in both mutants (0.52 ± 0.025, 0.67 ± 0.035, 0.5 ± 0.019, 0.48 ± 0.018 for control, Doc2b^WT,DN,6A^). Together, these data suggest that both mutants increase the initial release probability and impair prolonged neuronal secretion.

### Doc2b^DN^or Doc2b^6A^mutants have no effect on synaptogenesis

Doc2b is temporarily and spatially regulated during the embryonic and early postnatal phase (Korteweg et al., 2000) suggesting a role in neuronal development and synaptogenesis. In addition, Syt-7, another high affinity Ca^2+^-sensor which shares homology with Doc2b, has been implicated in neurite outgrowth (Arantes and Andrews, 2006). One plausible explanation, for the high frequency of spontaneous release in Doc2b^DN^and Doc2b^6A^expressing cells would be that those mutations affect neurogenesis or development and increase the synaptic density of neurons via altered membrane trafficking. To evaluate this possibility, we performed immunostainings for the synaptic vesicle marker Synaptophysin and the dendritic neuronal marker microtubule associated protein 2 (MAP-2, Figure 6). Quantitative morphometry of autaptic neurons expressing Doc2b^WT^and mutants did not reveal significant changes in synaptic density, dendritic length, dendritic synapse density, synapse area soma area or synapse distance from the soma (Figure 6B). Thus, the spontaneous release rise induced by Doc2b mutants is not caused by developmental dysregulation.

**Figure 6.**
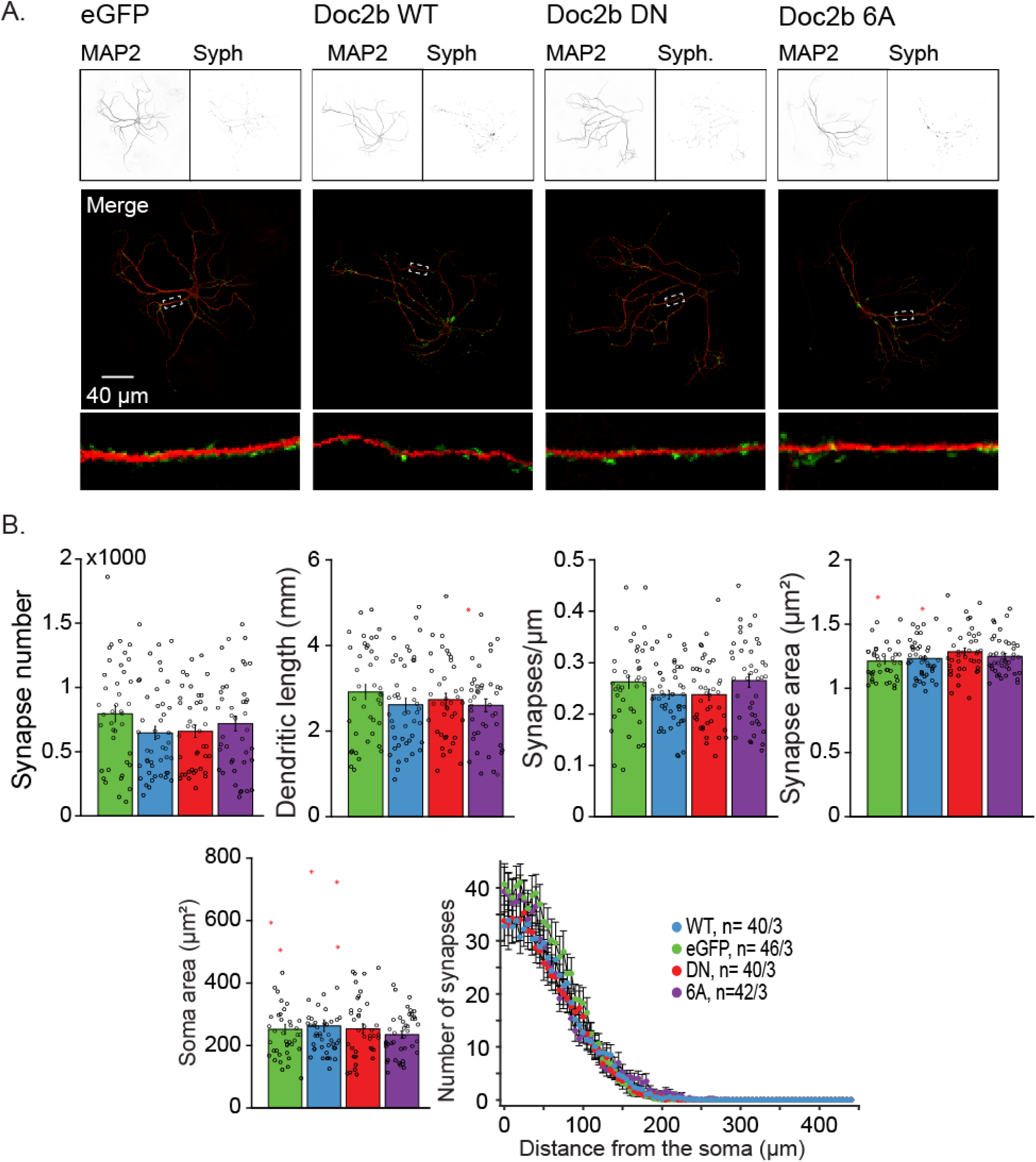
Overexpression of Doc2b^WT^, Doc2b^DN^, and Doc2b^6A^does not affect morphology or synapse number in wildtype neurons. (**A**) Confocal images from autaptic wildtype neurons expressing Doc2b constructs or control (eGFP). MAP2 (top left) immunostaining reveals dendritic morphology of the neurons. Synaptophysin (Syph; top right) stains for presynaptic active zone. White squares on merged images depict zoomed in dendritic areas (bottom). (**B**) Quantification of synapses and morphologic characteristics (Schmitz et al., 2011). Data are represented as mean ±SEM from the indicated number of cells over 3 independent experiments. Kruskall Wallis ANOVA and one way repeated ANOVA, Tukey’s Post-hoc tests.

### Doc2b^DN^or Doc2b^6A^mutants enhance spontaneous release in Doc2-deficient neurons

To rule out possible effects of endogenous Doc2b in our experiments, we investigated release in Doc2a/b double knock-out (DKO) autaptic neurons (Figure 7). The neurons were infected with Doc2b^WT^, Doc2b^DN^or Doc2b^6A^and expression was confirmed by immunoblotting (Figure S1B). Expression of Doc2b^WT^increased the average mini frequency from 10 ± 1.4 to 13 ± 1.6 Hz in DKO cells. This trend is similar to previous observations (Groffen et al., 2010) but was not significant in this case, likely due to the smaller sample size and overall high mini frequencies in this experiment. Both mutants again caused a strong increase in the mEPSC frequency while the amplitude (Figure 7C), rise and decay (Figure S2C) were unaffected. Quantification of the mEPSC charge showed a small but significant increase for Doc2b^6A^expressing neurons compared to other groups (Figure S2C, effect size (r): 0.838, Supplemental Table 1). Again, the effect size was low. Taken together, the high mEPSC frequency upon Doc2b^DN^and Doc2b^6A^expression is replicated in DKO neurons, further confirming a gain-of-function effect of these mutations in resting neurons.

**Figure 7.**
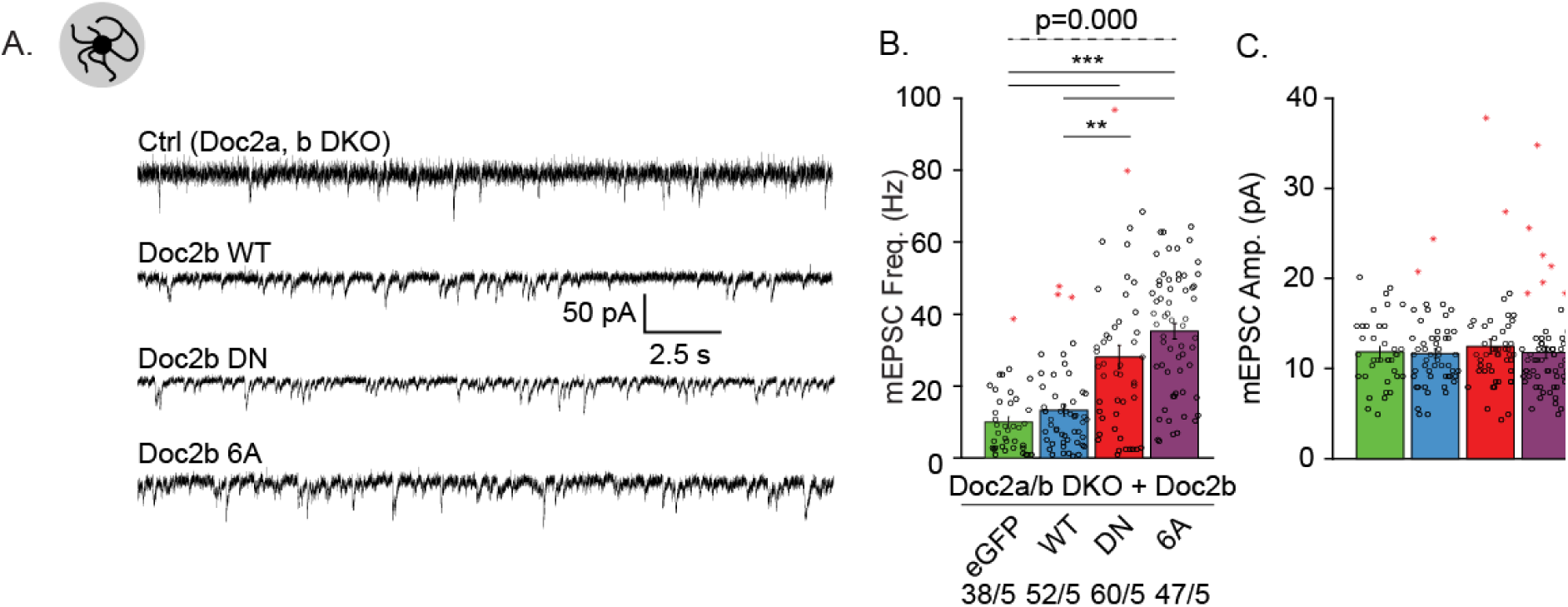
Effect of Ca^2+^binding site mutants of Doc2b on spontaneous release in Doc2-deficient neurons. (**A**) Representative mEPSC recording in autaptic neurons from Doc2a & b knockout mice after overexpression of eGFP (ctrl), Doc2b^WT^, Doc2b^DN^and Doc2b^6A^. (**B**) Quantification of spontaneous neurotransmitter release frequency and charge. Data are represented as mean ±SEM Kruskall Wallis ANOVA, Pairwise Post-hoc tests (*, p<0.05, ***, p<0.005). Red stars signs indicate outliers (outliers were included in the analysis)

### Faster synaptic depression in DKO neurons expressing mutant Doc2b^DN^or ^6A^

Similar to the results in wildtype cells, expression of Doc2b^WT^in DKO cells did not affect the 1^st^evoked EPSC charge (Figure 8A-B). The 1^st^evoked EPSC charge in Doc2b^DN^and Doc2b^6A^mutant expressing cells showed the same tendency to increase (Figure 8B) as in WT neurons, although this increase was not significant. Both mutants caused stronger depression during paired pulse stimulation (Figure 8C), again most notably in the 20-200ms interval range. Faster depression also occurred during repetitive stimulation at 5 and 40 Hz (Figure 8E,H,K,N), again without affecting the overall neurotransmitter release. The balance in synchronous and asynchronous release was slightly shifted by Doc2b^WT^expression in favor of the asynchronous component during 5 Hz trains (Figure 8I), suggesting an effect of the wildtype protein on evoked asynchronous release. Again, Doc2b^WT^expressing cells showed a more complete recovery of synaptic strength 2 s after repetitive stimulation, an effect that was lacking in both Doc2b mutants (Figure 8P,Q).

**Figure 8.**
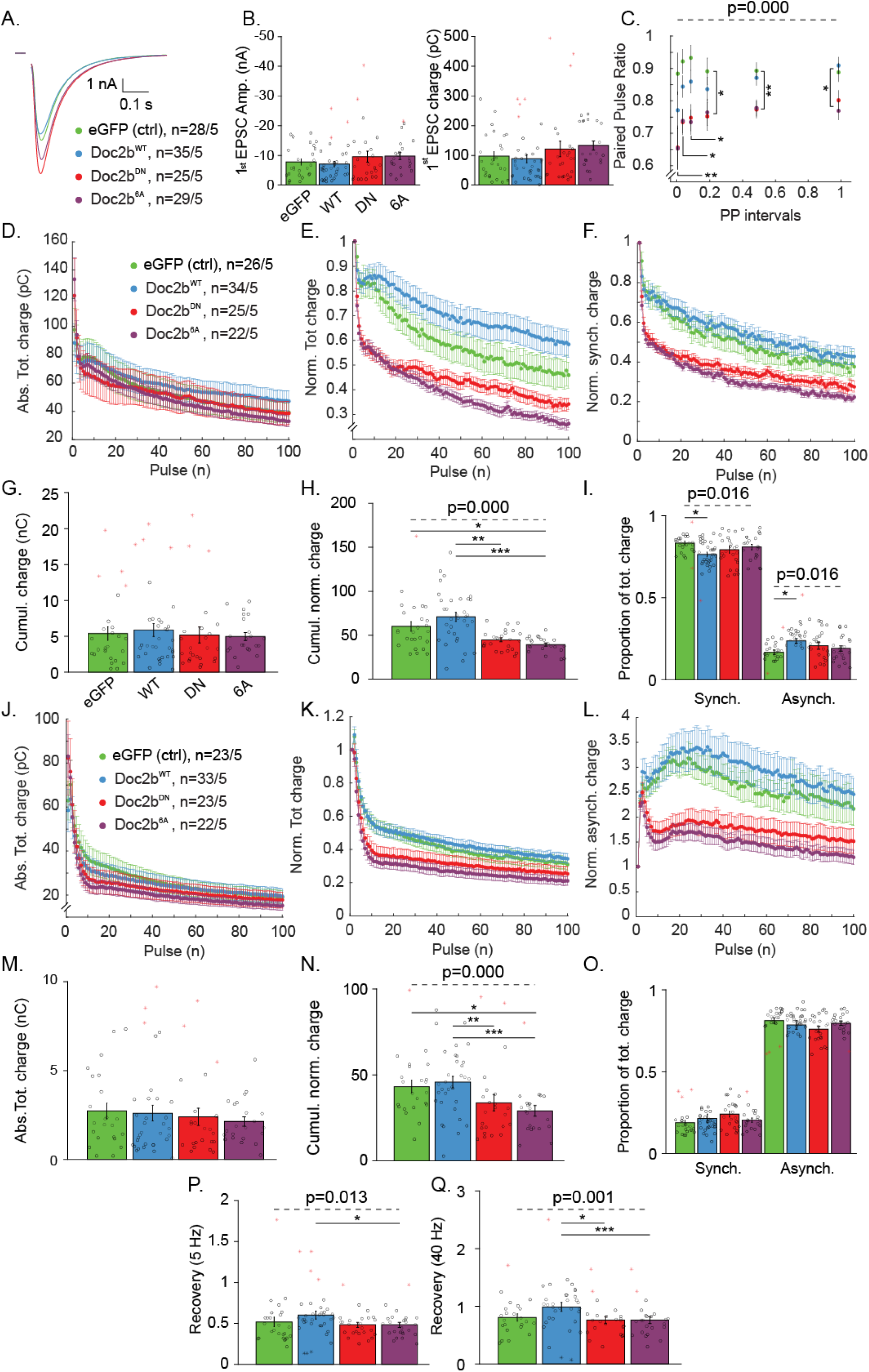
Ca^2+^binding site mutants Doc2b^DN^and Doc2b^6A^enhance short-term depression in Doc2-deficient neurons. (**A**) Average single EPSC from autaptic Doc2a & b knockout neurons expressing Doc2b^WT^and mutants. (**B**) Quantification of 1^st^EPSC amplitude and charge and (**C**) paired pulse ratio. (**D-I**) Rundown during repetitive stimulation at 5Hz. (**D**) Absolute and (**E**). normalized total charge rundown; (**F**) Normalized synchronous charge component. (**G**) Quantification of absolute and (**H**) normalized cumulated total charge transfer. (**I**) Proportion of synchronous and asynchronous charge components. (**J-O**) Rundown during repetitive stimulation at 40 Hz. (**J**) Absolute and (**K**) normalized total charge. (**L**) Normalized asynchronous release component. (**M**) Quantification of absolute and (**N**) normalized cumulative total charge transfer. (**O**) Proportion of synchronous and asynchronous charge components. (**P)** Recovery, measured as normalized EPSC charge after 100 stimulations at 5 Hz followed by 2 s rest. (**Q**) Recovery measured likewise after a 40Hz train. Data are represented as mean ±SEM Kruskall Wallis ANOVA, Pairwise Post-hoc tests (*, p<0.05, **, p<0.01, ***, p<0.005).

## Discussion

To shed light on Doc2b protein function in synaptic release, we studied the Ca^2+^binding site mutants Doc2b^DN^and Doc2b^6A^. We found that both mutants i) mimic an activated state at low [Ca^2+^] resulting in constitutive membrane enrichment and increased spontaneous neurotransmission; ii) have impaired Ca^2+^-induced membrane enrichment capacity and negatively affect synaptic strength during repetitive activity (paired pulse, train stimulation, recovery after repetitive stimulation). These phenotypes completely mirrored the phospholipid binding behavior in cell-free assays and cultured neurons, suggesting that phospholipid association is a key event in Doc2b’s overall function in secretion.

### The gain– or loss–of–function paradox

In previous studies, Doc2b^DN^has been interpreted as a gain-of-function mutant based on the increased phospholipid binding at rest (i.e. the mutations were considered to mimick Ca^2+^-binding). Doc2b^6A^was described to be a loss-of-function mutant based on the loss of Ca^2+^binding capacity. This classification as gain-and loss-of-function mutants had important implications: the enhanced spontaneous release in Doc2b^DN^overexpression was taken as evidence that Doc2b functions as a Ca^2+^sensor (Groffen et al., 2010), while the very similar phenotype in Doc2b^6A^overexpressing neurons was taken to demonstrate a Ca^2+^-independent function (Pang et al., 2011).

Our data show that Doc2b^6A^shows constitutive membrane binding, as supported by other studies (Houy et al., 2017; Courtney et al., 2018). Thus, Doc2b^6A^is not a pure loss-of-function mutant. At the same time however, the Doc2b^DN^mutant is also not a pure gain-of-function mutant, as indicated by several negative effects during repetitive neuronal activity. The dual effect of Ca^2+^binding site mutations could be attributed to the different surface charge distribution of the aspartates in the Ca^2+^-and membrane binding site of the C_2_ domains. In Doc2b wildtype protein, the aspartates residues are neutralized by binding of Ca^2+^ions. In the mutants, the neutralization of the aspartates could support membrane binding at rest, but this binding may not reach the same affinity as with Ca^2+^-bound aspartates. Alternatively, the mutations could slightly misplace the membrane-inserting residues in the loops surrounding the aspartates.

The Ca^2+^-dependent behavior membrane association of both mutants seems slightly more complex as might be expected. At high (>500 nM) [Ca^2+^]_free,_ isolated C_2_AB fragments showed a loss of liposome clustering activity using synthetic membranes composed of 75%DOPC and 25%DOPS. Recent investigations revealed a similar effect for the Doc2b^6A^mutant (termed 6x) as its liposome binding capacity dropped in presence of Ca^2+^ (Courtney et al., 2018). In our hands a similar reduction in lipid binding occurred at high (>700 nM) [Ca^2+^]_free_ for Doc2b^WT^C_2_AB domain. In live neurons where the physiological membrane composition is different, incomplete detachment was also observed for Doc2b^6A^(Figure 2C) but not for Doc2b^WT^or Doc2b^DN^. Syt-7 was also reported to have a reduced lipid binding activity at high [Ca^2+^]_free_, effect that was associated with an inhibitory effect on norepinephrine secretion in PC12 cells (Sugita et al., 2001). Another similar observation is the negative effect of high [Ca^2+^]_free_on Syt-1 in vitro fusion ability (Park et al., 2015), together suggesting that most Ca^2+^-sensors function in a limited [Ca^2+^] window.

In a previous study using isothermal calorimetry, the D218,220N mutation did not affect Ca^2+^ binding activity in a recombinant C_2_AB fragment named C_2_A_CLM_B (for calcium ligand mutant; Gaffaney et al., 2014). However, C_2_A_CLM_B_CLM_ carrying the additional mutation D357,359N in the C_2_B domain abolished Ca^2+^association to C_2_domains, suggesting that in the absence of lipids, the C_2_B domain is solely responsible for the Ca^2+^binding activity (Gaffaney et al., 2014). The C_2_A_CLM_B_CLM_mutant also greatly increased spontaneous release.

In presence of phospholipids, the negatively charged head groups may stabilize bound Ca^2+^ions, increasing the apparent Ca^2+^affinity (Zhang et al., 1998; Radhakrishnan et al., 2009). D220N substitution within the C_2_A Ca^2+^-binding-pocket of Doc2b, alone or in combination with other mutations is responsible for the constitutive membrane binding (Xue et al., 2015). A single residue substitution (D303N) completely abolishes Doc2b translocation and this mutant does not rescue spontaneous release (Courtney et al., 2018), confirming the idea that Doc2b acts as a Ca^2+^-sensor. Considering all these mutants, there is a striking correlation between the Ca^2+^-dependent phospholipid association of Doc2b and its function in spontaneous neurotransmission at rest.

### Ca^2+^ binding mutants affect synaptic plasticity during repetitive activity

In chromaffin granule secretion, wildtype Doc2b serves both positive and negative roles (Houy et al., 2017), a feature that is shared with other exocytotic proteins such as synaptotagmins, complexins and munc18s. In this system, Doc2b^DN^and Doc2b^6A^favour immediate chromaffin granule fusion but impair sustained release at high [Ca^2+^]. This phenotype might also be a collateral effect of constant vesicle fusion at rest, exhausting the immediate releasable pool (IRP)

In synapses, we observed a clear modification of evoked release by Ca^2+^binding mutants. However, overexpression of Doc2b^WT^(Figure 5 and 8) or the removal of endogenous Doc2b (Groffen et al., 2010) does not importantly affect evoked release, suggesting that this is not a major function of Doc2b in synapses. The effect of Doc2b mutants on evoked release may possibly represent an ectopic function. On the other hand, Doc2b^WT^overexpression and rescue revealed a consistent effect in post-burst recovery (Figure 5R,S and Figure 8P,Q), absent in DN and 6A expressing neurons. This could occur by increasing the vesicular release probability at high residual [Ca^2+^] after burst activity. Doc2b^DN^and ^6A^ineffectiveness in this potentiation mechanism might be the result of their loss of Ca^2+^-sensitivity. Additionally, a subtle but significant shift in favour of asynchronous release during repetitive stimulation appeared in DKO neurons rescued with Doc2b^WT^(Figure 8I) but not with mutants. During intense release, as suggested by the reduced in vitro lipid binding at high [Ca^2+^], Doc2b^WT^could inhibit fast exocytosis acting as a clamp of the fast fusion machinery, sparing the ready releaseable pool (RRP) and preserving a slowly releasable pool (SRP; Houy et al., 2017).

Mutant overexpression caused short-term depression during trains and paired pulse stimulation in both wildtype and DKO neurons. A higher 1^st^EPSC coupled to short-term depression in mutant expressing neurons suggests an increase in vesicle release probability (P_vr_) in the early phase of release. In autapses we found no evidence that Doc2b mutants enhance asynchronous release, as was previously observed for locally stimulated neuronal networks (Gaffaney et al., 2014; Xue et al., 2015). We speculate that an asynchronous release component could possibly build up from delayed synchronous release indirectly connected to the postsynaptic cell.

## Conclusion

We conclude that the Doc2b^DN^and Doc2b^6A^mutants do not represent divergent gain-and loss-of-function mutants but show similar behavior, characterized by increased activity at rest and impaired activity at high [Ca^2+^]_i_during neuronal activity. When using mutations to investigate the role of Doc2b in asynchronous or spontaneous release, it is important to consider the different performance of the protein at low and high Ca^2+^. The strict correlation between plasma membrane association and spontaneous release frequency supports a direct role as a Ca^2+^sensor. In addition, a Ca^2+^-dependent function in synaptic recovery is also supported by the data. Our collective results provide a unifying explanation for seemingly conflicting data and emphasize the importance of the Ca^2+^-dependent phospholipid association in Doc2b-mediated secretory regulation.

## Material & Methods

### Mouse lines

Animals were housed, bred and handled in accordance with Dutch and EU governmental guidelines. Protocols were approved by the VU University Animal Ethics and Welfare Committee (approval number FGA 11-06). Wildtype C57BL/6J mice were obtained from Charles River Laboratories. Doc2 a & b double knockout mice (DKO), maintained on the same C57BL/6J genetic background, were previously described (Groffen et al., 2010). To dissociate brain tissue from DKO mice, hippocampi were isolated at postnatal day 1 (P1). For wildtype mice, E18–stage embryos were used. In this case, pregnant females were sacrificed by cervical dislocation, embryos were obtained by caesarian section, decapitated and used for dissection.

### Primary culture of mouse neurons

To isolate mouse neurons, brains were placed in Hanks buffered salt solution (HBSS, Sigma) buffered with 1 mM HEPES (Invitrogen). After meninges removal, hippocampi and cortices were dissociated and separately treated. The tissue was incubated with 0.25% trypsin (Invitrogen) for 20 min at 37°C and washed in DMEM. Cells were dissociated by trituration with a fire-polished Pasteur pipette and counted in a Fuchs-.Rosenthal chamber. Neurons were plated in warmed Neurobasal medium supplemented with 2% B-27, 1.8% 1M HEPES, 0.25% glutamax and 0.1% Pen-strep (all products Invitrogen) as previously established (Wierda et al., 2007).

Electrophysiology experiments were performed in network or autaptic cultures. For network cultures, hippocampal neurons were plated at a density of 25K cells per well in 12-wells plates on etched glass coverslips containing a confluent layer of rat astrocytes (Wierda et al., 2007). For autaptic cultures, 1.5K cells per 12-well or 3K per 6-well were plated on coverslips with astrocyte micro-islands stamps (Wierda et al., 2007). For live imaging, coverslips were coated with rat tail collagen solution (BD Biosciences, Bedford, USA) and cells were plated in low density networks (1K per 12-well) (Wierda et al., 2007). For western blotting, cortical neurons were plated at 300K per well in 6-well plates without coverslips, coated overnight with 0.0005% poly-L-ornithine (Sigma) and 2 µg/ml laminin (Sigma) in PBS and washed with sterile water.

### Viral overexpression of Doc2b

For functional assays, Doc2b and EGFP were expressed as separate proteins from a single mRNA using an IRES2 internal ribosome entry site. Wildtype rat Doc2b^WT^(LIP#1984) was compared to Doc2b^DN^carrying the D218, 220N mutation (LIP#1985) (Groffen et al., 2004, 2010) and Doc2b^6A^carrying the D163, 218, 220, 303, 357, 359A mutation (LIP#1986) (Pang et al., 2011). Lentiviral infectious particles were packaged in HEK293T human embryonic kidney cells with a passage number lower than 25, maintained in Dulbecco’s Modified Eagle Medium (DMEM) supplemented with 10% fetal calf serum, 50 U/ml penicillin-streptomycin and 1x non-essential amino acids (Gibco). At 2 days in vitro (DIV2) the cells were transfected at 50% confluence with three plasmids: p.MDG2 (encoding the viral envelope protein), pCMVδR8.2 (encoding packaging factors) and a p156RRL-derived plasmid encoding Doc2b. LIP#1984 encoded Doc2b^WT^, LIP#1985 Doc2b^DN^and LIP#1986 Doc2b^6A^. At DIV3 the medium was changed to Optimem + 50 U/ml penicillin-streptomycin without fetal bovine serum. At DIV4, the supernatant containing infectious particles was centrifuged at 1000 x g to remove cell debris. The supernatant was concentrated by ultrafiltration using a 100 kDa cutoff membrane (UFC910024, Millipore, spun at 4000 x g for 20-30 min) to achieve a final volume of 150 µl. The LIPs were diluted to 1 ml with phosphate-buffered saline, filtered through 0.45 µm and stored in aliquots at −80°C until use. Neurons were infected at DIV1 to induce Doc2b expression. To investigate subcellular protein localization, Doc2b was C-terminally tagged with EGFP. Neurons were transduced with Semliki infectious particles 10 to 12 hours before experimentation (SIP#293 encoding Doc2b^WT^, SIP#244 encoding Doc2b^DN^and SIP#295 encoding Doc2b^6A^) as described (Houy et al., 2017).

### Electrophysiology in primary hippocampal neuronal networks and autapses

For electrophysiology, both continental and island cells were used between DIV 14 to 21. Doc2b-expressing cells were identified by monitoring EGFP fluorescence. The standard extracellular medium included 140 mM NaCl, 2.4 mM KCl, 4 mM CaCl_2_, 4 mM MgCl_2_, 10 mM HEPES, 10 mM glucose, 300 mOsm, pH 7.3. Our standard intracellular (patch pipette) solution was EGTA free to prevent Ca^2+^buffering; it constituted 125 mM K-gluconate, 10 mM NaCl, 4.6 mM MgCl, 4 mM K_2_-ATP, 15 mM creatine phosphate and 10 U/ml phosphocreatine kinase, 300 mOsm, pH 7.3. To record spontaneous excitatory events in network cultures, 1 µM tetrodotoxin (TTX, Abcam) and 20 µM gabazine (Sigma Aldrich) were added to the extracellular medium. In autapses, only gabazine was added. Where indicated, the Ca^2+^ionophore calcimycin (A23187, Sigma) was used at a final concentration of 10 µM and applied by puff for 100 seconds.

The patch pipettes were made of borosilicate and pulled using a multi-step filament pulling (P-1000, Sutter Instruments, Novato, USA) to achieve a pipette resistance of 3 to 5 MOhm. In whole-cell configuration, neurons were voltage clamped at −70 mV with an Axopatch 200B or Multiclamp 700B amplifier (Molecular Devices). Signal was low-pass filtered at 1 kHz and digitized at 10 kHz with a Digidata 1440A or 1550 (Molecular Devices). Neurons with a series resistance (Rs) exceeding 15 MOhm or with an Rs increase beyond 20% of the initial value were excluded. Rs was compensated to 70%. EPSCs were elicited by depolarizing the cell to 0 mV for 1 ms. Standard stimulation paradigms comprised spontaneous activity recording, paired pulse stimuli with intervals from 20 ms to 1 s, two trains of each 100 action potentials at 5 Hz and 40Hz. Each train was followed by a single stimulus at 2s after the last depolarization to test synaptic recovery.

Miniature EPSCs (mEPSCs) were detected using Mini Analysis 6.0 (Synaptosoft Inc.), using thresholds of 7 pA for event amplitude and 15 pC for area. Evoked release events (paired pulse stimulations) were analyzed using an in-house routine in the MATLAB^®^environment (He et al., 2017 Mathworks) to calculate the paired pulse ratio, the EPSCs charge and amplitude. In AP-induced burst EPSCs, synchronous and asynchronous components were isolated from total response charge as follows. Stimuli artefacts were removed, synchronous charge was determined by a straight line between the EPSC starting point n and the following EPSC starting point n+1. The area below this line was considered as synchronous, the area between the line and before stimulation baseline was considered as asynchronous (see Figure 5K). Both components were calculated using cubic interpolation. Calcimycin evoked responses were quantitated using Clampfit 10.4 (Molecular Devices) by measuring the total charge transfer during the compound application.

### Doc2b live microscopy

For Doc2b protein translocation imaging, hippocampal neurons from wildtype mouse at embryonal day E18 were dissociated and platted at 25K per well on glia layer. Cells were double-infected with Semliki virus encoding for Doc2b^WT^-EGFP and either Doc2b^DN^-mCherry or Doc2b^6A^-mCherry between DIV9 and DIV11. The two SIP stocks were first mixed in a 1:1 volume ratio and then added to each coverslip. Live imaging was performed 8–11 h post infection using a Nikon A1R confocal laser microscope controlled by NIS-elements AR software version 4.30 (Laboratory Imaging). Intracellular and extracellular solutions were similar to electrophysiology experiments. Field stimulation of 15 seconds at 40Hz was given after 15 seconds of baseline recording. Trains were triggered from a Master 8 connected to a stimulus isolator (WPI type A385) set to 30 mA output current. ImageJ was used for data analysis. Doc2b translocation was measured by plotting line profiles going through the soma of the neuron.

### Solutions for phospholipid-binding assays

Chelated Ca^2+^/EGTA solutions containing 50 mM HEPES, pH 7.4, 100mM KCl, and varying concentrations of Na_2_EGTA and CaCl_2_(0 to 10mM) were made by predicting [Ca^2+^]free with MaxChelator (http://maxchelator.stanford.edu/CaEGTA-TS.htm). Actual [Ca^2+^]_free_ was verified in each solution by recording fluorescence excitation spectra of fura-2 (Invitrogen, 0.07 µM) at an emission wavelength of 510 nm on a LS55 fluorescence spectrophotometer (Perkin Elmer). [Ca^2+^]_free_ was calculated as Kd x [(R-R_min_)/(R_max_-R)] x (F_max_^380^/F_min_^380^), where F^380^is the fluorescence intensity at ʎ_excit_. = 380 nm and R is the ratio F^340^/F^380^. Kd_EGTA_ was measured at 34.906 nM. Liposomes were formed by drying chloroform solutions containing 25% 1,2-dioleoyl-*sn-*glycero-3-phospho-L-serine (DOPS, Avanti Polar Lipids) and 75% 1,2-dioleoyl-*sn*-glycero-3-phosphocholine (DOPC, Avanti Polar Lipids)under a nitrogen stream. The phospholipids were resuspended in 50 mM HEPES, 100 mM KCl, pH 7.4 to a final concentration of 1 mg/ml, sonicated 5 times for 10 seconds and centrifuged for 90 min at 21,000 x g to clear the liposomes from large aggregates as described previously (Friedrich et al., 2008).

### Expression and purification of Doc2b C_2_A and C_2_AB fragments

The C_2_A fragment (aa125-255) and C_2_AB (115-412) fragment of Doc2b^WT^, Doc2b^DN^and Doc2b^6A^were expressed as glutathione-S-transferase (GST) fusion proteins in the E. coli strain BL21 and purified as described (Brouwer et al., 2015). GST-C_2_A was eluted from glutathione-agarose beads by glutathione, leaving the GST tag attached and allowing GST dimerization. C_2_AB was eluted with thrombin digestion, thus removing the tag. In both cases, Ca^2+^-dependent C_2_-membrane interaction causes liposome aggregation which can be measured by an OD 350 nm increase (Connell et al., 2008a; Friedrich et al., 2008). Protein amounts, potential contamination or degradation were verified by SDS gel electrophoresis. To compare lipid-binding activities, recombinant proteins were pooled from the following number of expression cultures: 4 for C_2_A^WT^, 5 for C_2_A^DN^, 5 for C_2_A^6A^, 3 for C_2_AB^WT^, 6 for C_2_AB^DN^, 6 for C_2_AB^6A^.

### Phospholipid-binding assays

To measure phospholipid binding, 20 µl of liposomes were mixed with 78, 70, 70 µl for GST-C_2_A^WT^, GST-C_2_A^DN^, GST-C_2_A^3A^respectively and 73, 70, 60 µl for C_2_AB^WT^, C_2_AB^DN^, C_2_AB^6A^respectively of buffered Ca^2+^/EGTA solution in a quartz cuvette to a final concentration of 0.5 mg/ml and the absorption at 350 nm was monitored for 10 minutes at 0.2 s intervals in a Cary 50 UV-Vis spectrophotometer. GST-C_2_A and C_2_AB protein used concentration were determined by measurement of their lipids aggregation capacity to reach a maximal OD 350nm around 0.5 and measured afterward by SDS-PAGE. After 60s of baseline recording, GST-C_2_A or C_2_AB protein was added to a final concentration of 9 µM, 26 µM, 24 µM for GST-C_2_A^WT^, GST-C_2_A^DN^, GST-C_2_A^3A^respectively and 0.36 µM, 0.39 µM, 0.9 µM C_2_AB^WT^, C_2_AB^DN^, C_2_AB^6A^respectively, inducing liposome binding and consequently an increase of A_350_. The A_350_ increase was not observed in absence of Ca^2+^ (Figure 3).The EC50 values were manually calculated using OD 350nm half-maximum X-intercept from raw data.

### Immunostaining and confocal imaging for synapse counting

After lentiviral expression of Hippocampal neurons expressing Doc2b^WT^, Doc2b^DN^or Doc2b^6A^, as marked by IRES-eGFP fluorescence, were fixed for 20 minutes at RT in 3.7% paraformaldehyde. After washing with PBS, cells were permeated with 0.5% Triton X-100 for 5 minutes and incubated for 20 minutes with 2% normal goat serum and 0.1% Triton X-100 to prevent aspecific binding. Coverslips were incubated for 2 hours at RT or overnight at 4°C in presence of polyclonal chicken anti-MAP2 (Abcam, ab5392) and polyclonal guinea pig anti-Synaptophysin1 (SySy, 101004) both diluted 1000-fold. After washing, cells were incubated overnight at 4°C with Alexa-546-and 647-conjugated secondary antibodies (1:1000, Invitrogen), washed again and mounted with Mowiol. Images were acquired with a confocal microscope LSM 510 (Carl Zeiss) with 488 nm, 543 nm, 633 nm lasers, using a 40x oil immersion objective and a scan resolution of 1024 x 1024 pixels. Stacks of images with an optical thickness of 0.4 µm were obtained. Neuronal morphology characteristics were analyzed with an automated image analysis MATLAB^®^(Mathworks) routine (Schmitz et al., 2011).

### Western blots

Cortical neuron cultures from WT and Doc2a, b DKO mice were plated to 200 K cells per well, infected at DIV 1 with lentiviral Doc2b constructs (LIP #1984, #1985, #1986 as above) and harvested at DIV 17. Cells were washed 2 times with PBS, lysed in Laemmli sample buffer, loaded with 50% or the totality of each well for WT and DKO cells respectively, separated by SDS-PAGE and blotted on PVDF membrane (Biorad). Membranes were blocked in 2% skim milk powder (Merck) and 0.5% FCS (Gibco) in PBS with 0.01% Tween-20 (Sigma-Aldrich). Doc2B polyclonal antibody 13.2 was used as primary antibody (1:500) for incubation overnight at 4°C. Goat anti rabbit alkaline phosphatase (Jackson lab) was used as secondary and Attophos (Promega) as substrate for 30 min incubation at RT. Reprobing was made for actin immunostaining with monoclonal anti-actin antibody C4 (1:3000 Chemicon) and Goat anti mouse alkaline phosphatase as secondary Ab (1:10000).

### Statistical analysis

All statistical analysis was performed using SPSS v.25.00 (IBM Corp., Armonk, NY, USA). All data are reported as mean ±SEM, except when specified. The number of measurements “n”, indicate the number of cells per group and the number of independent observations “N”, represent experimental weeks. Data were checked for normality using Shapiro-Wilk and Kolmogorov-Smirnov tests. Homogeneity of the variance was assessed with Levene’s test and the sphericity assumption was tested with Mauchly’s test. Parametric or non-parametric tests were run depending on homogeneity assumption respect or violation. Moreover, if the sphericity assumption was not met the Greenhouse-Geisser correction was used to adjust the degree of freedom. Considering that groups were independently acquired from one another, independent samples tests were performed i) For two groups a Mann-Whitney U test was performed ii) For more than 2 groups, one-way repeated measure ANOVA or Kruskall-Wallis test were used. For each experiment, the p-value alpha significance threshold was adjusted for multiple testing. The effect size was calculated for the independent samples Mann-Whitney U test as 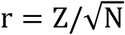, for one-way repeatedANOVA as 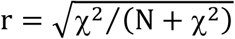 and for Kruskall-Wallis test as 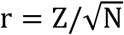 P-values lower than the accepted α-significance were highlighted in bold and the effect size was reported. Post-hoc tests were all pairwise comparison. In graphs, the empty dots represent single data points, bar plots represent the mean and error bars the SEM. Statistical outliers are depicted by red ‘stars’ symbols.

## Acknowledgements

We thank Robbert Zalm, Desiree Schut, Joke Wortel, Joost Hoetjes, Ingrid Saarloos and Eline Kompanje for excellent technical support. We express gratitude to Vincent Huson, Javier Emerador Melero, Rocio Diez Arazola and members of CNCR lab for their critical advice and their scientific support.

## Author contributions

Quentin Bourgeois-Jaarsma, Formal analysis, Investigation, Writing-review and editing; Matthijs Verhage, Conceptualization, Writing-review and editing; Alexander J. Groffen, Formal analysis, Funding acquisition, Investigation, Writing-review and editing

---

**Figure S1.**
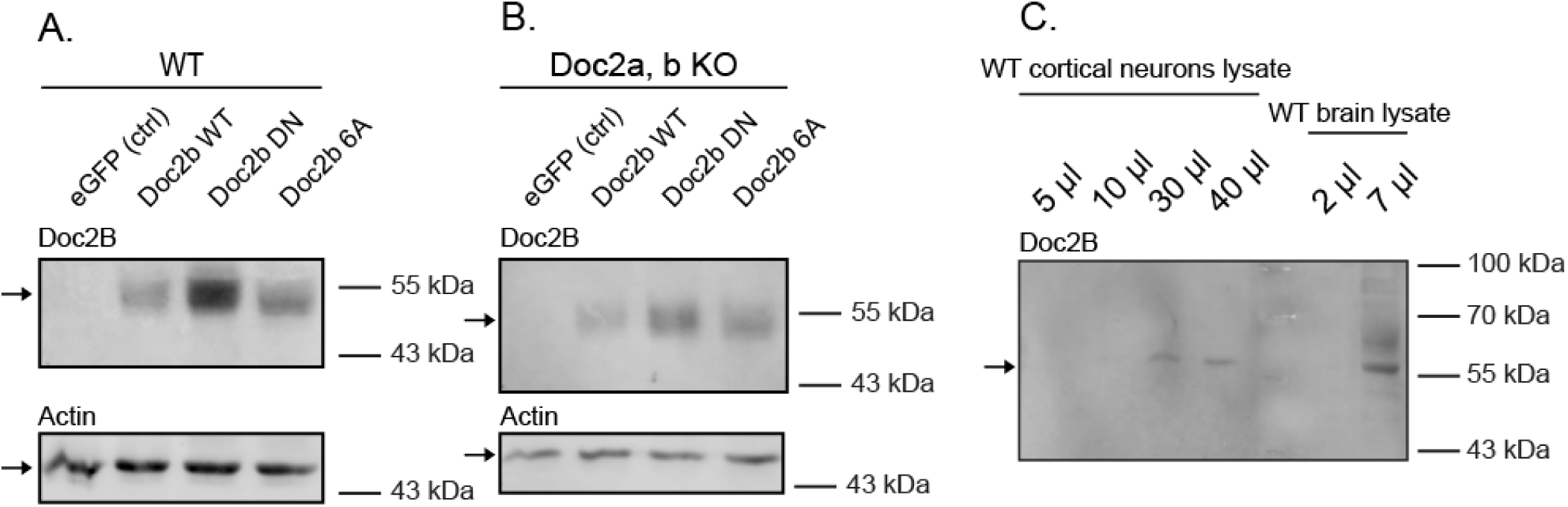
Doc2b expression levels in wildtype and Doc2a,b double knockout (DKO) neurons. Western blots from (**A**) wildtype and (**B**) DKO high density neuron cultures overexpressing wildtype or mutant Doc2b. (**C**) Endogenous Doc2b could not be detected in standard conditions but became detectable by loading larger sample volumes of cortex or total brain lysate. Doc2b immunoreactivity in Doc2 overexpressing neurons was higher compared to endogenous levels in non-transfected wildtype neurons.

**Figure S2.**
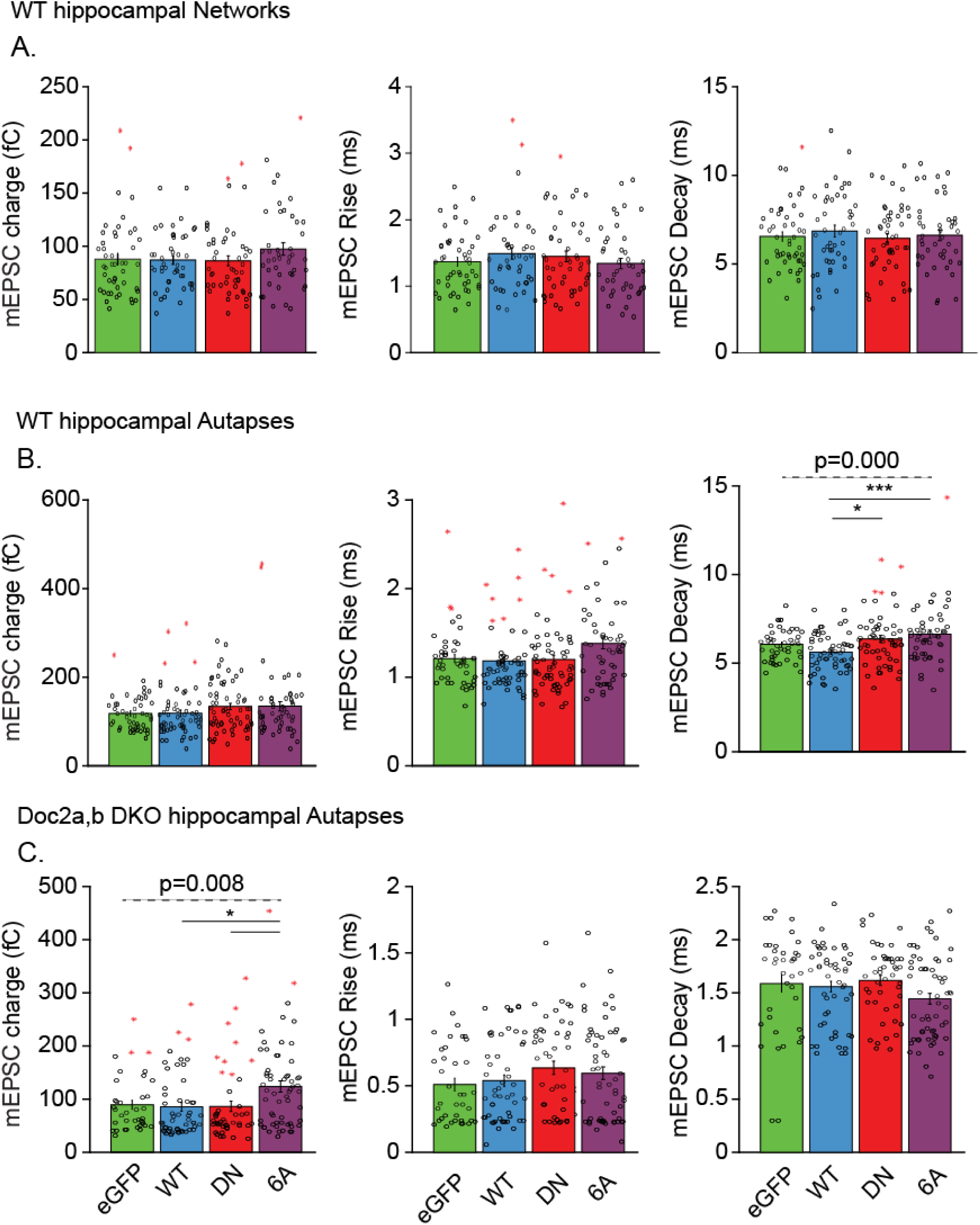
Postsynaptic characteristics of mEPSC events in cultured hippocampal neurons. (**A**) mEPSC amplitude, rise time and decay time in network cultures from wildtype mice; (**B**) same in autaptic cultures from wildtype mice; (**C**) same in autaptic cultures from Doc2-deficient mice. Data are represented as mean ±SEM.

**Figure S3.**
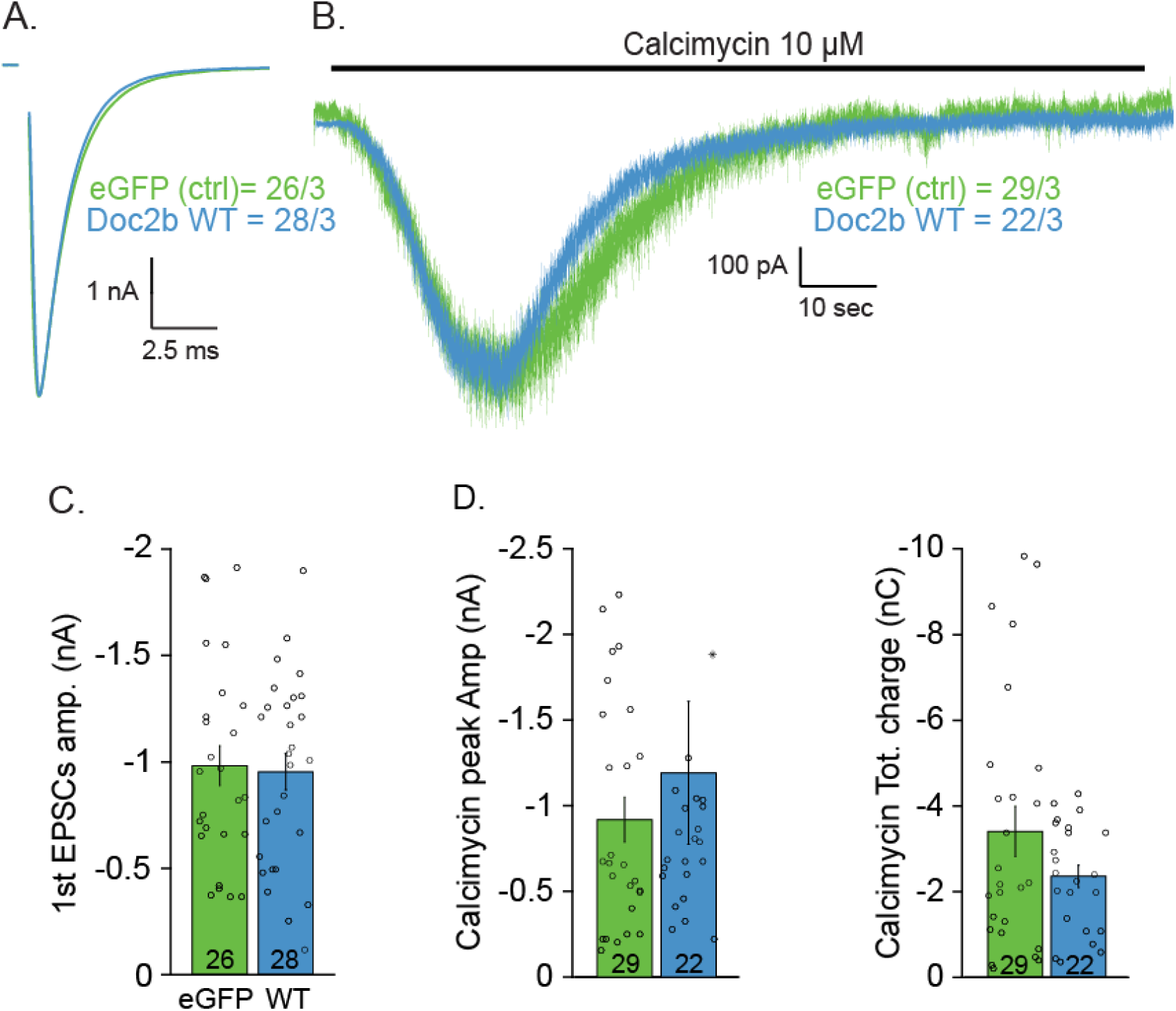
Overexpression of wildtype Doc2b does not affect calcimycin-induced EPSC charge. Calcimycin was perfused to directly induce Ca^2+^influx and bypass voltage-dependent Ca^2+^channels. (**A**) Typical example of single EPSC in naïve wildtype autaptic neurons; (**B**) the EPSC induced by puff application of calcimycin for 100 s. (**C**) Quantification of EPSC amplitude and calcimycin-induced charge transfer. Data are represented as mean ±SEM.

**Supplementary Table 1:**
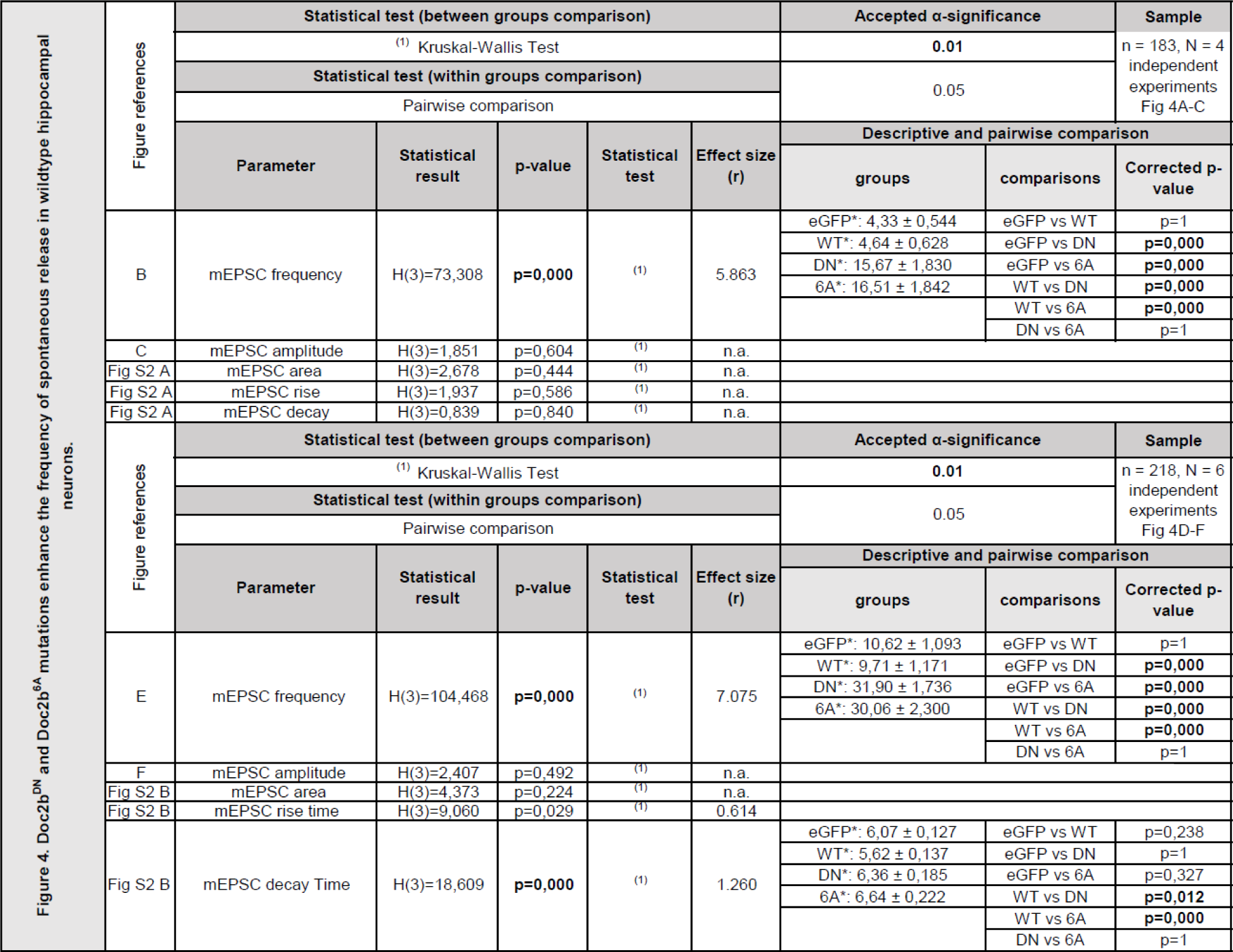
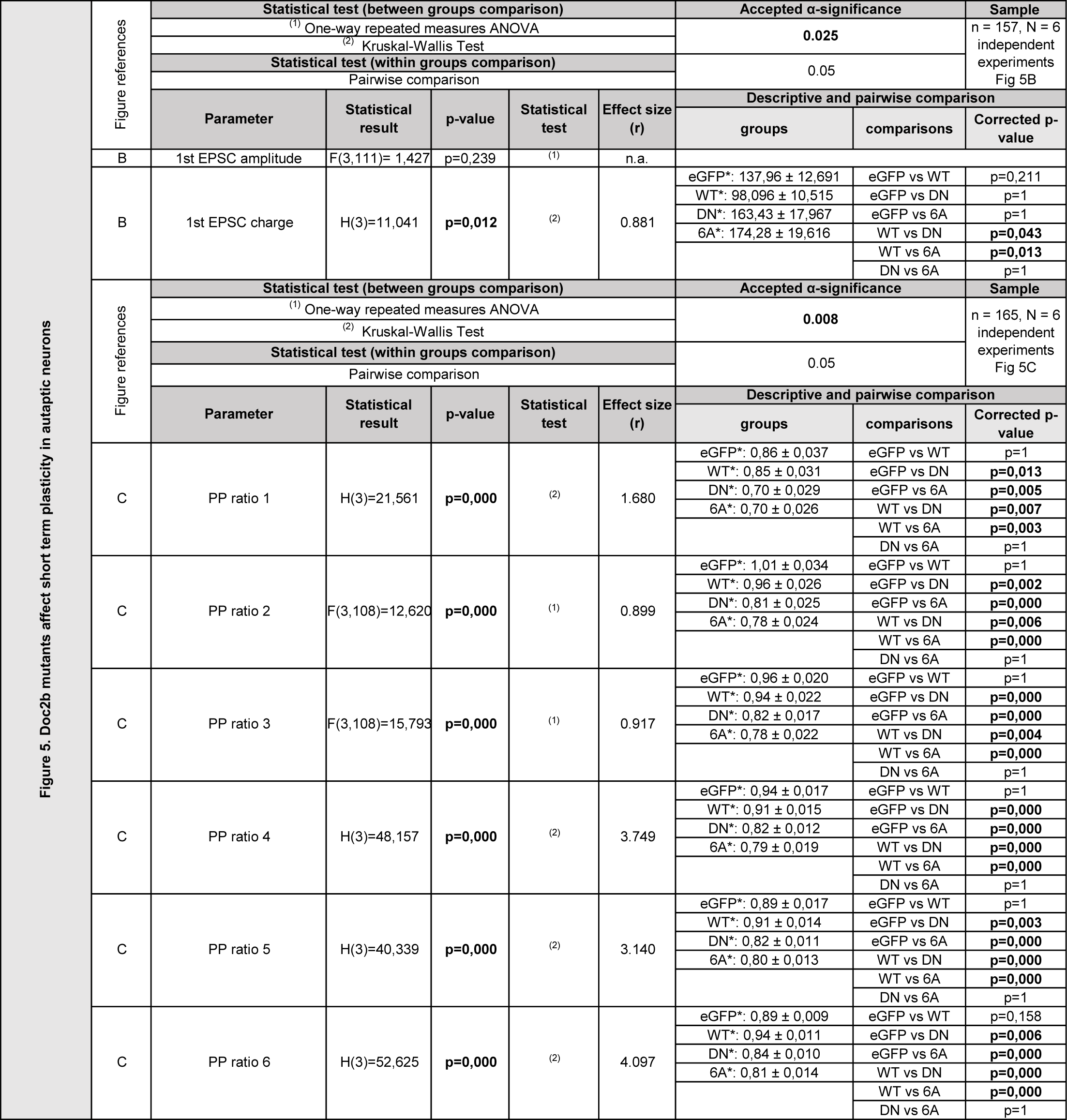
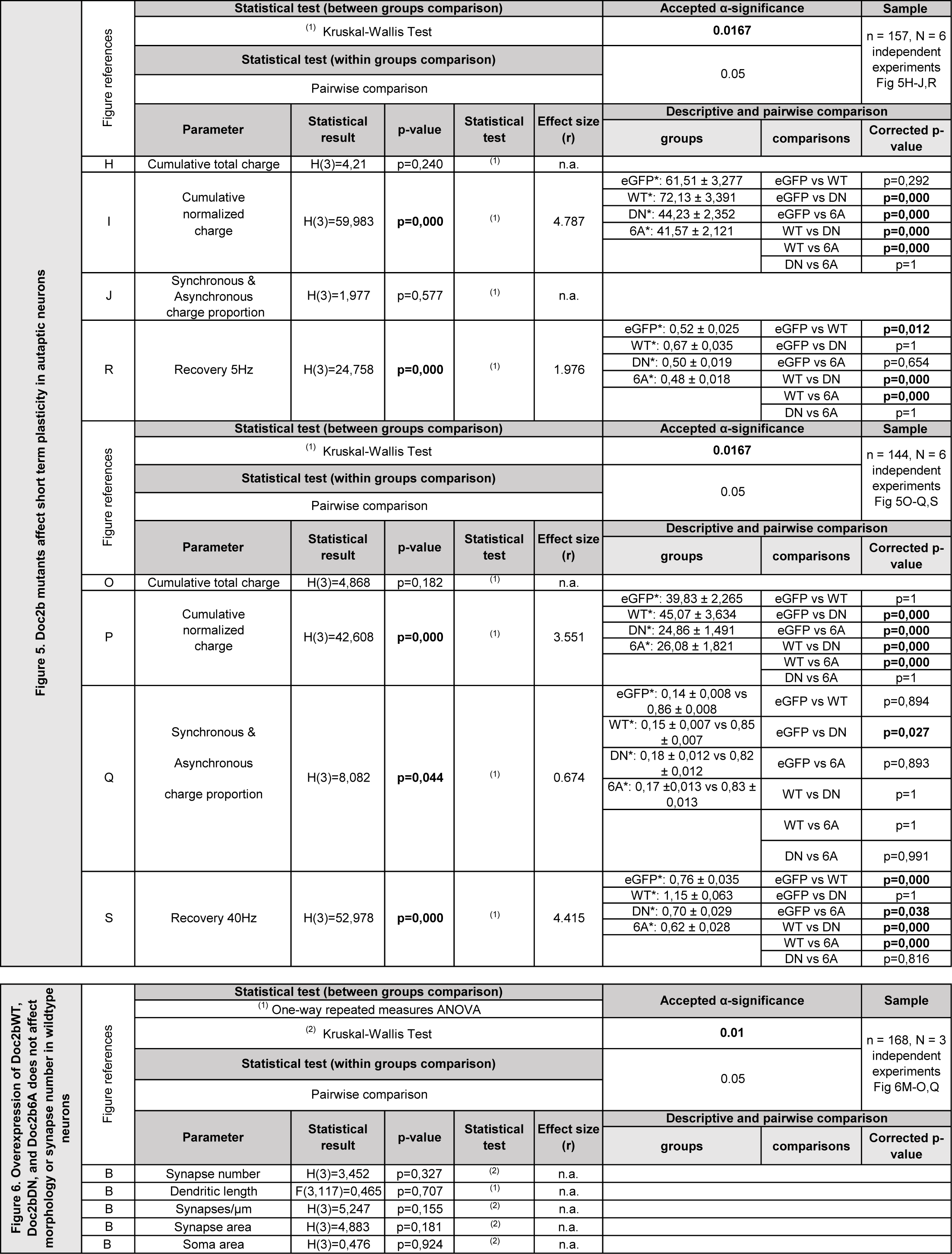
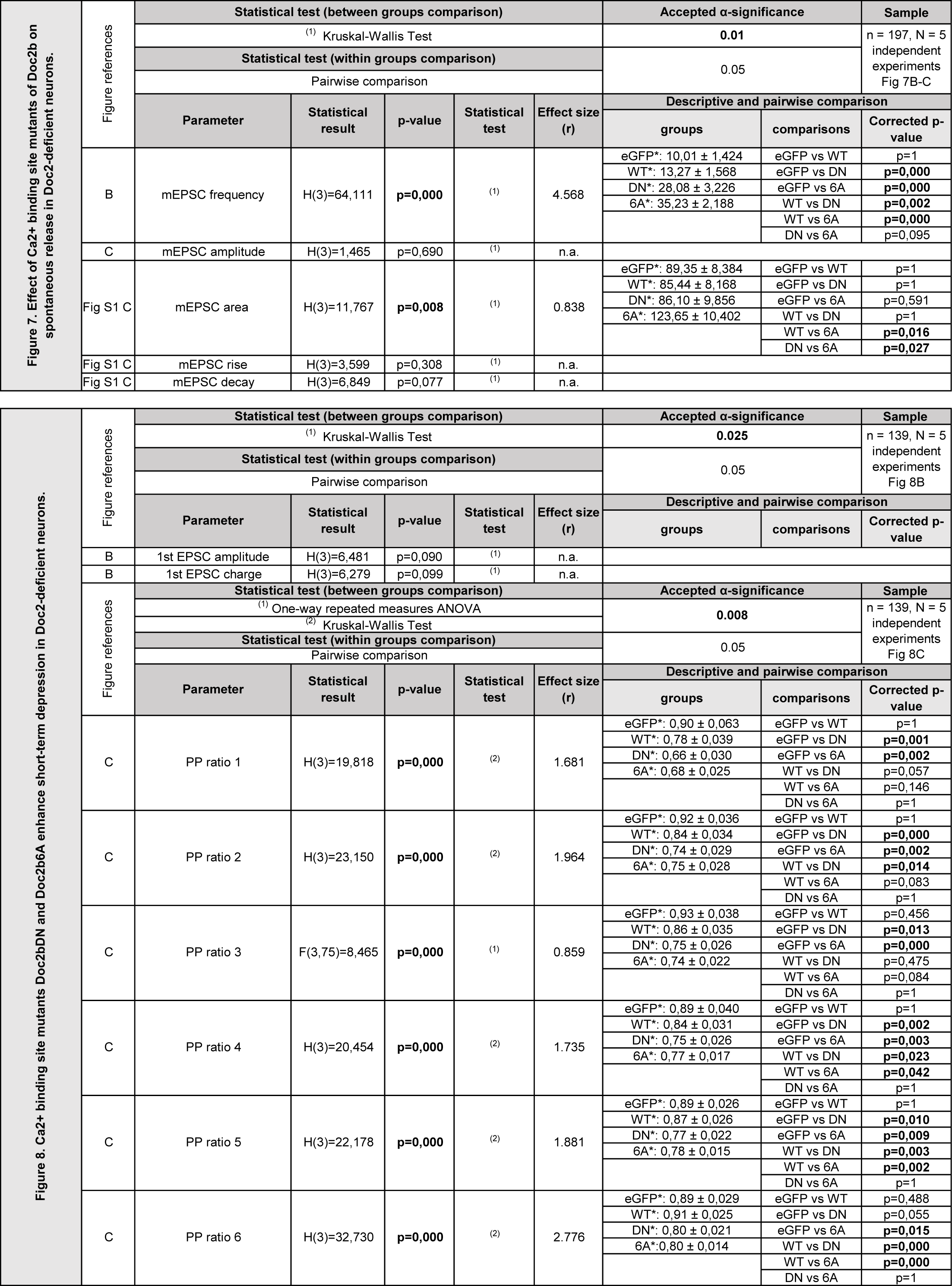
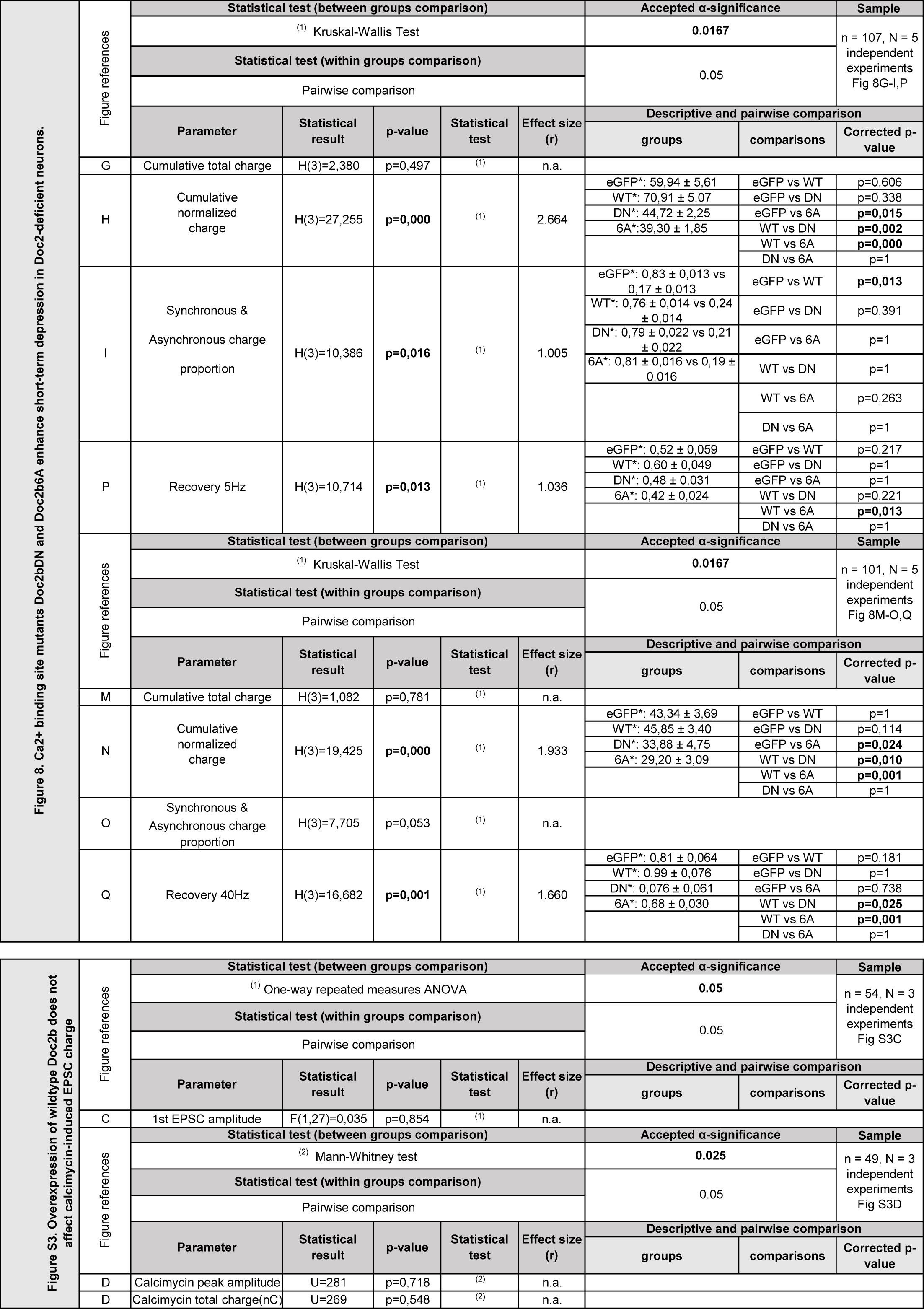
Statistics

